# Perturbing H-NS function reveals roles in restricting virulence heterogeneity and pathogen adaptation

**DOI:** 10.64898/2026.01.01.697318

**Authors:** Leah McLelland, Dakshya Karki, Madison Spratt, Miracle Burt, Jonathan Zhao, Keara Lane

## Abstract

Bacterial pathogens must balance rapid expression of virulence genes in host niches with tight repression when not needed to avoid fitness costs and ensure survival. Integration of virulence gene regulatory networks with the conserved global repressor H-NS was critical to achieve this balance. H-NS-mediated repression of virulence genes in non-inducing environments is essential for maintaining virulence genes and has shaped pathogen evolution. However, the role of H-NS-mediated repression in virulence gene activation and pathogen evolution in virulence-inducing conditions is less clear. For instance, although virulence gene expression is often heterogeneous, whether relief of H-NS repression contributes to this heterogeneity remains unknown. Furthermore, whether H-NS repression shapes pathogen evolution in virulence-inducing environments is unclear. Here, we use a *Salmonella* strain with reduced H-NS DNA-binding affinity to investigate the role of H-NS in virulence gene expression in individual bacteria and pathogen adaptation. We find that reduced H-NS repression increases the fraction of virulence gene expressing cells without eliminating bimodality, resulting in enhanced epithelial cell infection *in vitro*. Using experimental evolution, we demonstrate that the *hns* genotype constrains adaptive mutations and that disabling virulence gene expression is a common path to improved fitness in intracellular-like environments. Our results expand the role of H-NS-mediated repression from silencing virulence genes in non-inducing conditions to regulating heterogeneity in inducing conditions and demonstrate that evolutionary conservation of H-NS constrains adaptive strategies in intracellular-like environments.

## INTRODUCTION

Bacterial pathogens must rapidly adjust virulence gene expression patterns while colonizing diverse host environments [1–4]. Optimal virulence gene repression is a critical yet challenging part of this process. Excessive repression of virulence genes delays necessary functional responses, such as production of a type III secretion system (T3SS) or toxins, leading to a failure to adapt and survive in a host environment. Conversely, insufficient repression leads to inappropriate virulence gene expression, imposing fitness costs ranging from reduced growth to detection by the immune system [5–7]. To balance these competing demands, bacterial pathogens have evolved regulatory strategies that integrate virulence gene regulatory networks with global repressors, enabling both tight silencing and rapid activation as needed [8–10]. Here, we examine how perturbing one such global repressor, histone-like nucleoid structuring protein (H-NS), affects virulence gene expression heterogeneity and evolutionary adaptation in the intracellular bacterial pathogen *Salmonella enterica* serovar Typhimurium (STm).

H-NS is a global repressor of gene expression in Gram-negative bacteria, including STm [10,11]. It is an abundant nucleoid-associated protein composed of an N-terminal oligomerization domain and a C-terminal DNA-binding domain (DBD) connected via a flexible linker. The DBD preferentially binds to curved AT-rich DNA sequences, while the oligomerization domain enables higher-order complex formation that compacts and bridges DNA [12–17]. By binding AT-rich promoters, including horizontally-acquired virulence genes, H-NS occludes RNA polymerase and thereby blocks transcription [16,18].

H-NS-mediated repression is essential for STm viability. Loss of *hns* causes widespread misregulation of gene expression and significant growth defects, with Δ*hns* strains viable only in combination with mutations in genes such as *rpoS* (the stationary-phase sigma factor) or *phoP* (a two-component system response regulator) [16,18–20]. This contrasts with non-pathogenic *Escherichia coli* (*E. coli*), where *hns* is non-essential, implicating the misexpression of virulence genes as the basis for *hns* essentiality in STm [21,22]. Multiple virulence regulons are derepressed in STm Δ*hns* strains, and deletion of the virulence regulator *ssrA* partially rescues the growth defect of the Δ*hns* strain, demonstrating the contribution of H-NS-mediated repression of virulence genes to STm fitness [16]. While H-NS ensures tight repression of virulence genes under non-inducing conditions, this repression must be rapidly relieved in host environments. Host environmental cues, such as changes in osmolarity, pH, and ion concentrations, activate response regulators that displace H-NS from the promoters of virulence genes, enabling RNA polymerase recruitment and transcription [23–25]. Therefore, optimal virulence gene regulation depends on the coordinated interplay between H-NS-mediated repression and response regulator activity.

The interaction between H-NS-mediated repression and response regulator activity in virulence gene expression has been characterized primarily at the population level. However, single-cell studies reveal that virulence gene expression is highly heterogeneous in STm, even under uniform inducing conditions. This non-genetic variation, referred to as phenotypic heterogeneity, arises primarily from stochastic gene expression [26,27]. Generating distinct subpopulations of virulence-expressing and non-expressing bacteria enables bet-hedging or division-of-labor strategies that improve survival in hostile host environments [7,28]. In STm, phenotypic heterogeneity is associated with two major virulence regulons encoded in *Salmonella* Pathogenicity Islands 1 and 2 (SPI-1 and SPI-2) [29]. Expression of SPI-1, which is required for invasion of the intestinal epithelium, is bistable and incurs a fitness cost [5,30–32]. Both SPI-1(+) and SPI-1(−) subpopulations are required for a productive infection *in vivo*, and perturbing this ratio impairs infection outcomes. Expression of SPI-2, which is required for survival and replication within host cells, is bimodal and probabilistically tuned to the strength of intracellular-like signals, including pH and magnesium concentration, *in vitro* [31,33–38]. SPI-2 expression is heterogeneous within macrophages, with SPI-2(+) cells associated with anti-inflammatory gene expression profiles [39]. While the significance of phenotypic heterogeneity in STm virulence gene expression is well established, whether the relief of H-NS-mediated repression of virulence genes contributes to this cell-to-cell variation remains unknown.

While the role of H-NS in virulence gene expression heterogeneity remains unclear, it is well established that H-NS shapes pathogen evolution through its role as a global regulator of virulence gene expression [11,40,41]. H-NS-mediated repression is essential for maintaining virulence genes in the STm genome. Loss of *hns* causes growth defects that can be compensated by mutations in SPI-1 or SPI-2 genes, demonstrating that H-NS-mediated repression prevents fitness costs from the misexpression of virulence genes [16,40]. While the integration of virulence gene regulation with H-NS-mediated repression has enabled STm to maintain virulence genes and colonize diverse host environments, it may also act to constrain adaptive evolution. As a global regulator controlling the expression of hundreds of genes, H-NS may limit which genetic changes are beneficial as STm adapts to host environments. Understanding how H-NS impacts pathogen adaptation to intracellular environments may reveal how evolutionary constraints on this conserved global repressor shaped STm evolution in this critical host niche.

Here, we identify an H-NS DBD mutant strain of STm and use it to investigate how this global repressor regulates heterogeneity in virulence gene expression and constrains adaptation to intracellular-like conditions *in vitro*. By measuring virulence gene expression in individual bacteria, we find that reduced H-NS repression increases the fraction of virulence gene expressing cells without eliminating bimodality, and that this enhances epithelial cell infection. Using experimental evolution, we demonstrate that while both wild-type and mutant populations converge on disrupting virulence gene regulation to improve fitness in intracellular-like conditions, the *hns* genotype constrains the mutational paths available for adaptation. These results suggest that H-NS contributes to optimal virulence gene repression by setting activation thresholds in individual cells, and that strong purifying selection on this global repressor limits evolutionary adaptive strategies available to STm under intracellular-like conditions.

## RESULTS

### Identification of a *Salmonella* strain with a mutation in the H-NS DNA-binding domain

While investigating the growth of STm in SPI-2-inducing conditions (MgM-MES) *in vitro*, we identified a strain with impaired growth compared to the wild-type (WT), while its growth in non-inducing conditions (M9/Glc/CAA) was unaffected (Figure 1A). Quantification of growth metrics confirmed that this mutant (MUT) strain had a significantly increased lag time and a reduced maximum growth rate compared to the WT strain in MgM-MES (Figure 1A). To determine whether this impaired growth was generalizable to other virulence-inducing conditions, we cultured the strains in media used to induce the SPI-1 virulence regulon (LB High Salt: LB-HS) and found growth was comparable between the two strains (Figure S1A). MgM-MES not only induces SPI-2 but also imposes several stressors on the bacteria, including magnesium restriction, nutrient limitation, and acidic pH. To test whether stress conditions were sufficient to elicit a growth defect in the MUT strain, we measured growth in a nutrient-limited condition (M9/Glc without casamino acids) (Figure S1A). The MUT strain showed impaired growth in M9/Glc compared to the WT strain, with significant changes in lag time and maximum growth rate. Together, these data demonstrate that the MUT strain has context-specific growth defects, with impaired growth in SPI-2-inducing and nutrient-limiting media conditions.

**Figure 1.**
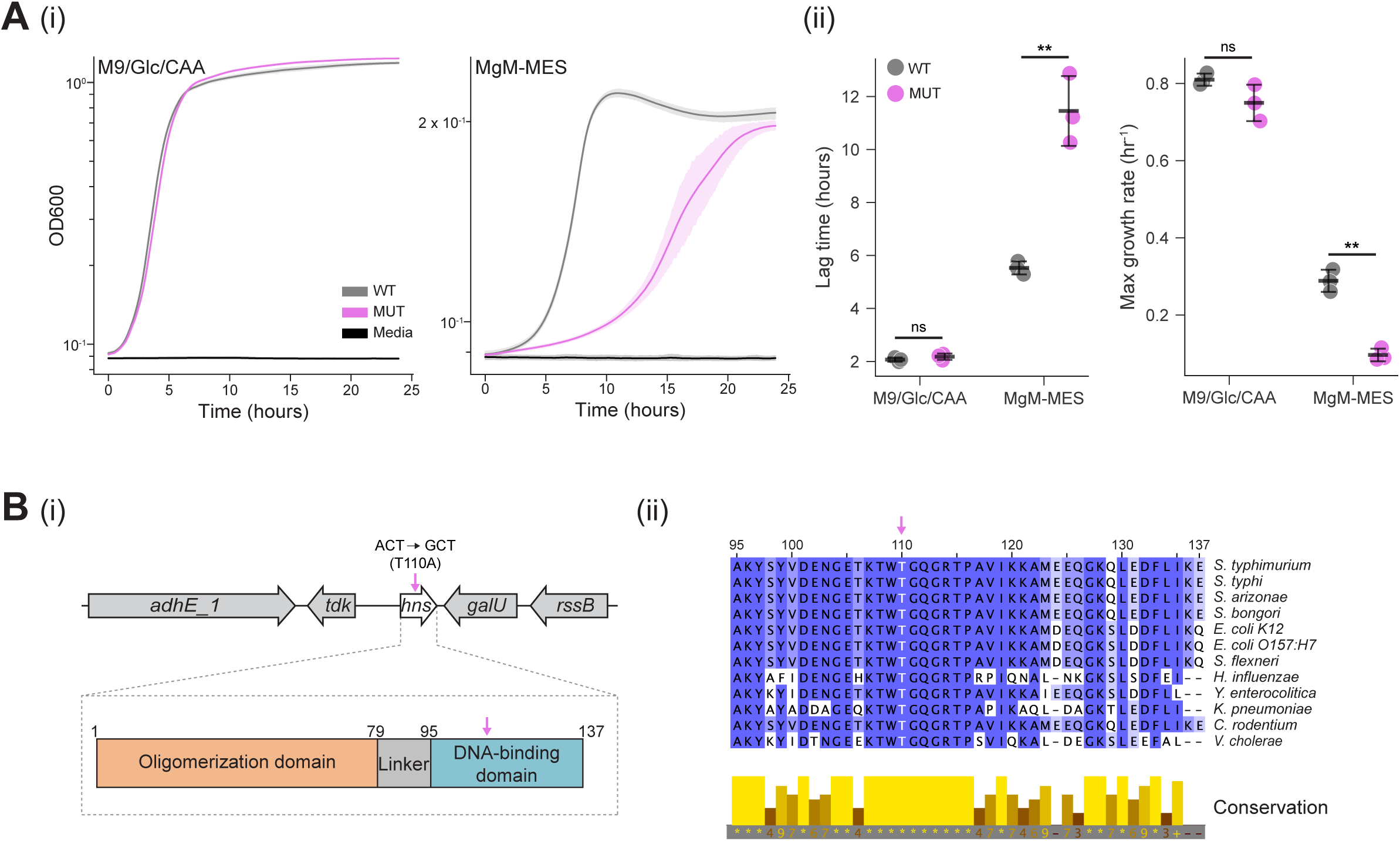
Identification of an H-NS DNA-binding domain mutation in a *Salmonella* strain with impaired growth in SPI-2-inducing conditions. (A) (i) The mutant strain exhibits growth defects in SPI-2-inducing conditions. Wildtype (WT, gray) and mutant (MUT, magenta) strains were grown in non-SPI-2-inducing media (M9/Glc/CAA) or SPI-2-inducing media (MgM-MES), and OD_600_ was recorded using a plate reader. Growth curves show averages of three independent experiments, each with three technical replicates; shaded regions represent standard deviation. Media control is shown in black. Y-axes are scaled to the growth condition. (ii) Growth metrics extracted from data in (i). Individual datapoints represent independent experiments. Asterisks denote significance as determined by two-sided unpaired t-test with Benjamini-Hochberg FDR correction as follows: ** p<0.01; ns, not significant. (B) The MUT strain contains a mutation in the conserved DNA-binding domain (DBD) of H-NS. (i) Schematic showing the genomic location of *hns* in STm and domain organization of the H-NS protein. Arrow indicates the location of the mutation identified by whole genome sequencing. (ii) Multiple sequence alignment showing conservation of the mutated residue (T110, white) across enteric bacteria. Dark blue and asterisks indicate complete conservation.

To identify the genetic basis for this context-specific growth defect, we performed whole genome sequencing on the WT and MUT strains. We identified a single missense mutation (a328g [T110A]) in the DNA-binding domain (DBD) of *hns* (Figures 1B, S1B). The H-NS DBD is highly conserved across enteric bacteria, with Thr110 completely conserved (Figure 1B) [42,43]. Several other genetic changes were identified in the MUT strain (Figure S1B, Table S1). However, these changes were located in either intergenic regions, a pseudogene, or in regions coding for hypothetical proteins, and were therefore considered unlikely to cause the impaired growth of the MUT strain. In STm, *hns* null mutations are viable only when accompanied by mutations in *rpoS* or *phoP* [16,18,20]; therefore, this suggests that T110A is a hypomorphic mutation that impairs but does not abolish H-NS repression. As such, the T110A strain is a unique tool to quantitatively assess how altered H-NS repression affects virulence gene expression in individual bacteria.

### H-NS T110A has reduced DNA-binding affinity

Based on the location of the T110A mutation in the highly conserved H-NS DNA-binding domain and the T110A strain’s growth defects, we hypothesized that the T110A mutation reduces H-NS DNA-binding affinity. To assess how the T110A mutation might alter protein structure, we used an NMR-resolved structure of the H-NS WT DBD from STm (PDB: 2L93) (Figure 2A) [15]. Two consequences of the T110A mutation that could negatively impact DNA binding were apparent. First, in the WT structure, Thr110 uses a side chain hydroxyl to form a polar interaction with Gln112, a residue in the QGR motif. This motif mediates direct contacts between H-NS and the DNA minor groove [15,44]. The substitution of Thr110 with alanine eliminates the side chain hydroxyl and replaces it with a methyl group, resulting in the loss of the stabilizing hydrogen bond between residues 110 and 112. Removing this polar interaction could destabilize the QGR motif and reduce H-NS DNA-binding affinity. Second, the T110A mutation alters van der Waals forces. In H-NS T110A, an interaction between Thr110 and Trp109 is lost, and a new interaction between Ala110 and Lys96 is gained. The Thr110 – Trp109 interaction likely anchors the polar bond between Trp109 and Gln112 to stabilize the QGR motif. This structural analysis confirms that Thr110 is a critical residue in the H-NS DBD and suggests that the T110A mutation disrupts key interactions in the DBD that are important for proper positioning of the QGR motif.

**Figure 2.**
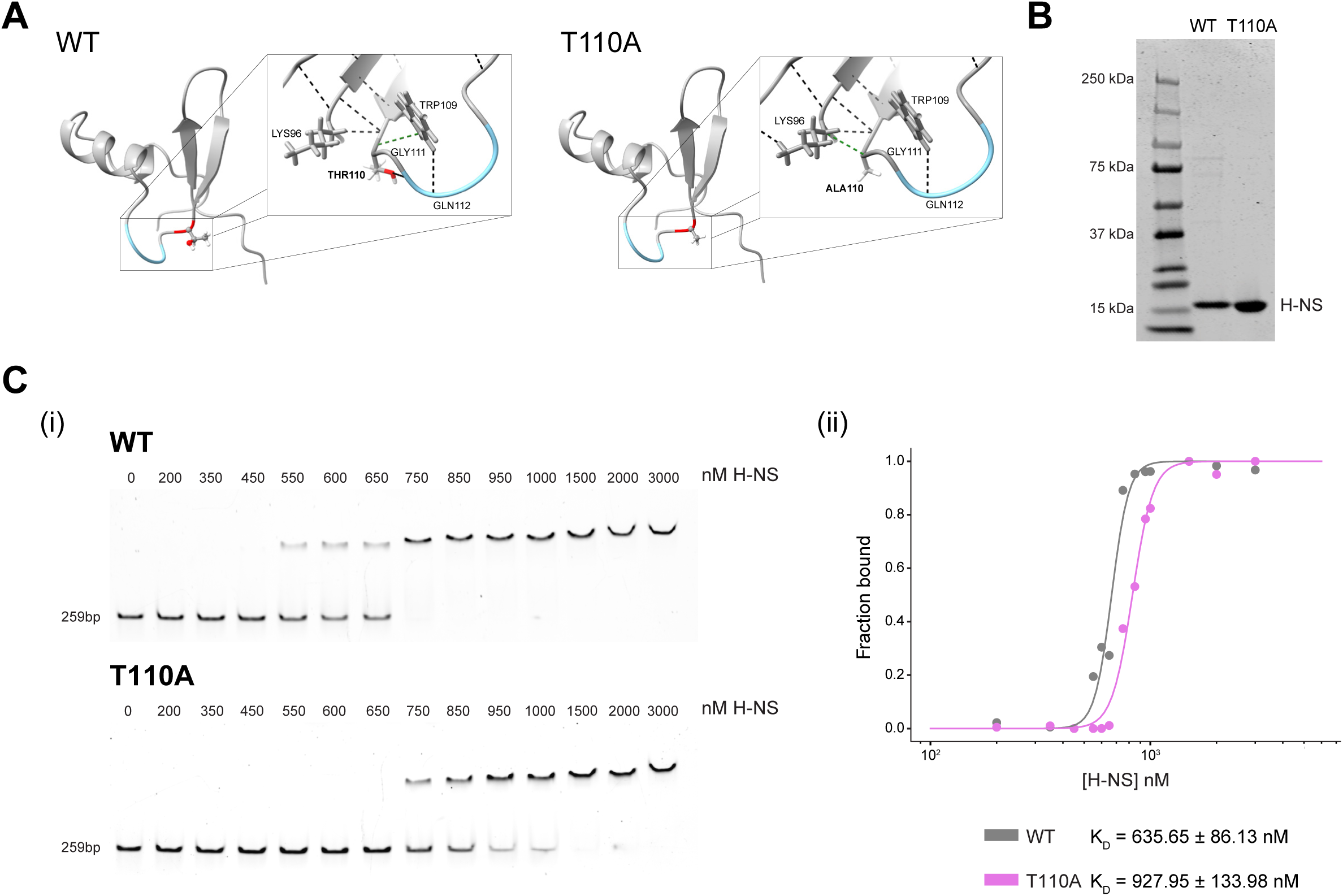
H-NS T110A has reduced DNA-binding affinity compared to H-NS WT. (A) Structure of STm H-NS WT and T110A DNA-binding domains modeled in ChimeraX using the NMR structure (PDB: 2L93). The T110A mutation is predicted to disrupt a polar contact (black dashed line) between the Thr110 side chain oxygen and the Gln112 backbone nitrogen. Gln112 is part of the QGR motif (shown in blue), an AT-hook that mediates direct DNA contacts. The mutation is also predicted to disrupt van der Waals interactions (green dashed line) between Thr110 and Trp109, while introducing a new interaction between Ala110 and Lys96. These structural changes may alter the positioning of the QGR motif and reduce its ability to bind the DNA minor groove. (B) Purified H-NS WT and T110A proteins analyzed by SDS-PAGE. Protein was visualized using Coomassie staining. (C) EMSAs demonstrate reduced DNA-binding affinity of H-NS T110A. (i) H-NS WT and T110A proteins were incubated with *ssrAB* promoter DNA at the indicated concentrations for 30 minutes and separated on a 5% native TBE gel. DNA was visualized using SYBR Safe staining. Representative gels are shown for WT (top) and T110A (bottom). (ii) Quantification of EMSA data from panel (i) showing the fraction of DNA-bound H-NS as a function of protein concentration. WT (gray) and T110A (magenta) data points are shown as circles and represent measurements from a single replicate. Lines represent fits to the Hill equation. K_D_ values shown are means ± standard deviation from four independent experiments.

Next, to test whether the T110A mutation reduces H-NS DNA-binding affinity, we performed electrophoretic mobility shift assays (EMSAs). We expressed and purified C-terminal 6xHis-tagged full-length H-NS WT and T110A in *E. coli* using polyethyleneimine (to remove DNA), nickel affinity chromatography, and size exclusion chromatography (Figures 2B, S2A-B). We selected a region in the *ssrAB* promoter as the DNA probe because *ssrAB* is part of the SPI-2 virulence regulon, is a known H-NS target, and this promoter region has been previously validated in H-NS EMSAs (Figure S2C) [24,45,46]. As predicted from the structural analysis, H-NS T110A retained DNA-binding capacity but had reduced affinity compared to H-NS WT (Figures 2C, S2D). Although there was some variation between replicates, the reduced affinity of H-NS T110A for DNA was reproducible. Free DNA probe was fully depleted between 750-950 nM for H-NS WT compared to 1500-2000 nM for H-NS T110A. Quantification of binding and fitting to the Hill equation confirmed that the average K_D_ for H-NS T110A (927.95 ± 133.98 nM) was higher than for H-NS WT (635.65 ± 86.13 nM) (Figures 2C, S2E). These results demonstrate that the T110A mutation impairs H-NS function by reducing DNA-binding affinity, consistent with findings for *E. coli* H-NS DBD mutants [47,48].

### H-NS T110A alters features of virulence gene expression heterogeneity while preserving bimodality

Virulence gene expression in *Salmonella* is heterogeneous, with only a subset of cells activating expression even under uniform virulence-inducing conditions [5,30–33,37,38]. H-NS plays a critical role in repressing virulence genes in non-inducing conditions [16,18]; however, the extent to which H-NS-mediated repression contributes to heterogeneity in virulence gene expression remains unclear. The T110A strain provided a tool to investigate this by allowing a direct comparison of single-cell virulence gene expression between strains with intact (WT) and impaired (T110A) H-NS-mediated repression.

Bimodality in virulence gene expression typically arises from stochastic gene expression and feedback in the gene regulatory network, generating subpopulations of responders and non-responders [49–52]. However, virulence gene expression also requires the cognate response regulators to displace H-NS from virulence gene promoters, allowing RNA polymerase recruitment and transcription [16,18,23–25]. We hypothesized that if H-NS-mediated repression contributes to heterogeneity in virulence gene expression, reduced H-NS DNA-binding affinity in the T110A strain could affect this heterogeneity in several distinct ways, which can be distinguished at the single-cell level (Figure 3A). For instance, in the T110A strain 1) the response could be faster if H-NS is displaced more easily, 2) the fraction of responders could increase if H-NS displacement is a limiting step in expression, or 3) the amplitude of expression could increase if H-NS rebinding limits transcriptional output. Comparing virulence gene expression between H-NS WT and T110A strains enabled us to determine whether H-NS-mediated repression contributes to heterogeneity in virulence gene expression.

**Figure 3.**
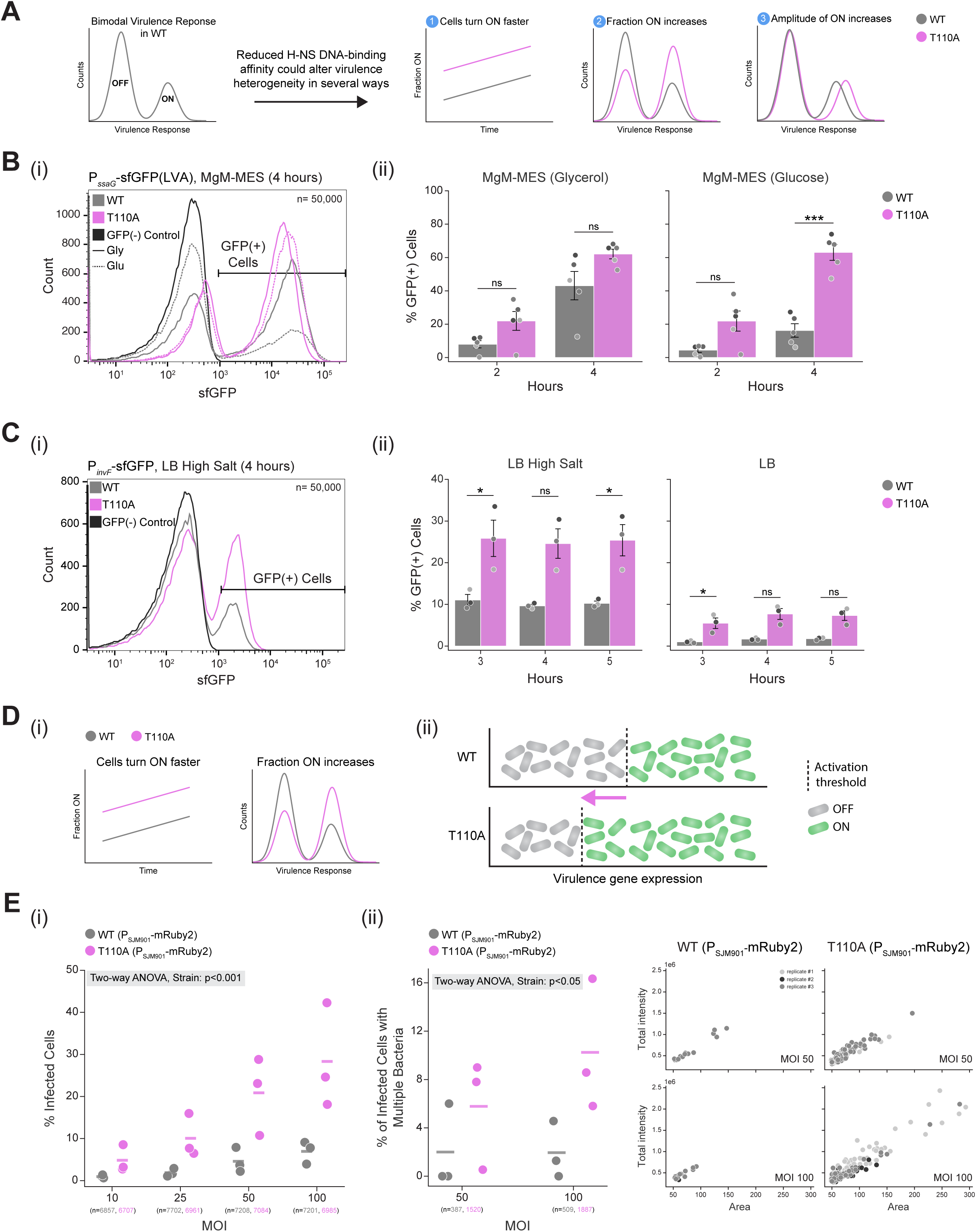
H-NS T110A increases the fraction of virulence-gene expressing cells with functional consequences for infection. (A) Schematic illustrating three potential outcomes of reduced H-NS DNA-binding affinity on virulence gene expression heterogeneity. (B) H-NS T110A cells activate the SPI-2 reporter more rapidly and with increased frequency, while maintaining bimodal expression. WT and T110A strains containing the SPI-2 reporter plasmid (P_ssaG_-sfGFP(LVA)) were grown in SPI-2-inducing media (MgM-MES) and analyzed by flow cytometry. (i) Representative histograms at 4 hours of WT (gray) and T110A (magenta) strains grown in MgM-MES with glycerol (solid line) or glucose (dashed line). GFP(−) control strain is shown in black. (ii) Quantification shows the percentage of GFP(+) cells at 2 and 4 hours. Bars represent the mean ± SD of 5 independent experiments. Individual datapoints are shown in shades of gray. n=50,000 cells per sample. (C) H-NS T110A increases the frequency of SPI-1 responders while maintaining bimodal expression. WT and T110A strains containing the SPI-1 reporter plasmid (P_invF_-sfGFP) were grown in SPI-1-inducing media (LB High Salt) and non-inducing media (LB) and analyzed by flow cytometry. (i) Representative histograms at 4 hours of WT (gray) and T110A (magenta) strains grown in LB High Salt. GFP(−) control strain is shown in black. (ii) Quantification shows the percentage of GFP(+) cells at 3, 4, and 5 hours. Bars represent the mean ± SD of 3 independent experiments. Individual datapoints are shown in shades of gray. n=50,000 cells per sample. For (B) and (C) asterisks denote significance as determined by linear mixed-effects model with post-hoc pairwise comparisons using unpaired t-tests with Holm-Sidak correction for multiple comparisons: * p<0.05, *** p<0.001; ns, not significant. (D) (i) Schematic illustrating features of virulence heterogeneity altered in the H-NS T110A strain. (ii) Activation threshold model for how H-NS alters the fraction of cells expressing virulence genes. In the T110A strain, the virulence activation threshold (dotted line) is lowered (pink arrow), increasing the responder fraction (green) while preserving bimodality. (E) H-NS T110A strain enhances epithelial cell infection. H-NS WT and T110A strains expressing constitutive mRuby2 (P_SJM901_-mRuby2) were used to infect HeLa cells at varying multiplicity of infection (MOI), and infections were imaged and quantified as described in the Methods. (i) Percentage of infected cells. Line represents the mean and circles the individual datapoints from 3 independent experiments. T110A results in an increased frequency of infected cells compared to WT (two-way ANOVA, effect of strain: F(1,16)=25.1, p<0.001; n=total cells across 3 independent experiments). (ii) Percentage of infected cells that contain multiple bacteria. Line represents the mean and circles the individual datapoints from 3 independent experiments. T110A shows an increased frequency of multi-bacteria infection events (two-way ANOVA, effect of strain: F(1,8)=6.4, p<0.05; n=total infected cells across 3 independent experiments). Scatter plot shows total intensity versus area of bacterial masks for infected cells containing multiple bacteria. Individual infected cells are represented as circles, and replicates are shown in shades of gray.

Given the impaired growth of the T110A strain in SPI-2-inducing media (MgM-MES, Figure 1A), we first examined whether the mutation altered SPI-2 virulence gene expression patterns. To measure SPI-2 transcriptional activity in individual bacteria, we used a previously described SPI-2 transcriptional reporter consisting of the *ssaG* promoter (a promoter upstream of the primary SPI-2 structural operon) fused upstream of a destabilized sfGFP (sfGFP-LVA) [33]. The reporter plasmid also constitutively expresses mRuby2 for cell identification by flow cytometry. In the WT strain grown in MgM-MES, SPI-2 reporter expression is bimodal, with bacteria progressively switching into the SPI-2 ON state over several hours [33]. We compared SPI-2 reporter expression between the WT and T110A strains in either SPI-2-inducing (MgM-MES (Glycerol)) or non-inducing (M9/Glc/CAA) media after 2 and 4 hours of growth. In SPI-2-inducing media, the T110A strain maintained bimodal SPI-2 reporter expression with distinct ON and OFF populations (Figure 3B). We next evaluated whether any of the three features of response heterogeneity (response timing, fraction of responders, and response amplitude) differed between the strains. To evaluate response timing and the fraction of responders, we compared the fraction of GFP(+) cells over time. At both 2 and 4 hours in SPI-2-inducing media, the T110A strain consistently had a greater fraction of responders than the WT strain (Figure 3B, Table S2, 5/5 replicates, sign test, p=0.0312). The increase in the fraction of GFP(+) cells at 2 hours, when cells first begin to express the SPI-2 reporter [33], indicates that SPI-2 reporter expression is faster in the T110A strain. To determine whether the increase in responder fraction over time in the T110A strain was a result of its slower growth rate in MgM-MES and protein accumulation, we substituted glucose for glycerol to increase the growth rate. At 4hrs, faster growth decreased the fraction of responders in the WT but not in the T110A strain, indicating that differences in growth rate do not fully explain the increase in responders in the T110A strain (Figure 3B, Figure S3A). Next, we evaluated the amplitude of the SPI-2 reporter response and found it was generally similar between the two strains, with only a modest increase in the mean fluorescence intensity (MFI) of the T110A GFP(−) subpopulation at 4 hours, which may result from slower growth (Figure S3B). As expected, in non-inducing conditions, the SPI-2 reporter remained OFF in both strains (Figure S3C), indicating that reduced H-NS DNA-binding affinity alone is insufficient to trigger reporter expression in the absence of activated response regulators. Together, these data indicate that reduced H-NS DNA-binding affinity in the T110A strain does not abolish the bimodal SPI-2 response but increases the response rate and the fraction of responders.

Unlike SPI-2, SPI-1 expression imposes clear fitness costs, including a reduced growth rate that is associated with increased antibiotic resistance [5,30]. In addition, the ratio of SPI-1(+) to SPI-1(−) cells is a critical feature of the system *in vivo*. Mutations that decrease the fraction of SPI-1(−) cells lead to the emergence of genetically avirulent cells and limit infection [32]. The fitness cost of SPI-1 expression together with the constraint on the responder fraction suggests that SPI-1 expression may be particularly sensitive to perturbations in H-NS-mediated repression. We therefore hypothesized that reduced H-NS DNA-binding affinity in the T110A strain would increase the fraction of SPI-1(+) cells.

To test this, we generated a SPI-1 transcriptional reporter consisting of the *invF* promoter (a SPI-1 response regulator) fused upstream of sfGFP, with constitutive mRuby2 expression for cell identification by flow cytometry. Similar plasmid-based reporters have been used in other studies to examine cell-to-cell variation in the SPI-1 response [53]. To test whether H-NS-mediated repression impacts the fraction of responders, we examined reporter expression in SPI-1-inducing conditions (LB High Salt) and quantified the fraction of responders at 3, 4, and 5 hours by flow cytometry (Figure 3C). The T110A strain maintained bimodal expression of the SPI-1 reporter but had a higher fraction of responders than the WT, and this difference remained stable over time (Figure 3C, Table S2). The MFI of the SPI-1(−) and SPI-1(+) subpopulations were similar between the strains (Figure S3D). As expected, in non-inducing media (LB), the SPI-1 reporter remained OFF in the WT strain (Figures 3C, S3E). In contrast, the T110A strain showed a small but consistent increase in the fraction of SPI-1(+) cells above the WT level that remained stable over time. Neither strain had reporter expression prior to the start of the assay, indicating this difference arose during the experimental growth period (Figure S3E). Since WT and T110A strains grow similarly in LB (Figure S1A), growth differences are not responsible for this increase in GFP(+) cells. It is possible that the use of a stable fluorescent protein results in GFP accumulation as cells enter stationary phase, potentially amplifying small differences in leaky expression caused by reduced H-NS repression in the T110A strain.

Both SPI-1 and SPI-2 responses remain robustly bimodal despite impaired H-NS function, demonstrating that H-NS does not determine whether virulence gene expression is heterogeneous. Instead, H-NS-mediated repression affects features of virulence heterogeneity, specifically the timing of the response and the fraction of responding cells (Figure 3D). The T110A mutation had a large effect on SPI-1 heterogeneity, suggesting that the activation threshold for SPI-1 virulence gene expression is sensitive to changes in H-NS DNA binding affinity (Figure 3D). In the T110A strain, lowering the SPI-1 activation threshold allows more cells to become SPI-1(+), while other known regulatory mechanisms maintain bimodality.

### Reduced H-NS function enhances *Salmonella* infection of epithelial cells

Expression of the SPI-1 regulon drives production of a T3SS required for invasion of epithelial cells [29,54,55]. Given the increased fraction of SPI-1(+) cells in the T110A strain, we asked whether this translates to enhanced epithelial cell infection. While the SPI-1 reporter reflects *invF* transcriptional activation, numerous subsequent steps are required for T3SS assembly and function [56]. In addition, a recent study of *E. coli* flagellar gene regulation revealed that distal genes can exhibit variable expression despite homogeneous expression of a proximal regulator [57]. As a result, the increased fraction of SPI-1 reporter ON cells in the T110A strain might not necessarily result in increased invasion capacity. Invasion can also be impacted by the ratio of SPI-1(+) to SPI-1(−) cells. The presence of both subpopulations was shown to maximize the invasion frequency of epithelial cells in culture, while a SPI-1(+) only subpopulation showed reduced invasion frequency [58]. To test whether the increased fraction of SPI-1(+) cells in the T110A strain increased epithelial cell infection, we infected HeLa cells with H-NS WT and T110A strains constitutively expressing mRuby2 and quantified infection using microscopy. We found the T110A strain increased the frequency of infected HeLa cells compared to WT across all multiplicities of infections (MOIs) tested (Figure 3E, Table S2). We further characterized infection by quantifying the fraction of infected HeLa cells containing multiple bacteria, using area and total intensity of segmented bacterial objects to distinguish between single and multi-bacteria infection events. At higher MOIs (50, 100), where multi-bacteria infection events are more likely to occur, the T110A strain showed an increased frequency of such events compared to the WT, as well as more bacteria per cell, as evidenced by greater area and total intensity of segmented bacteria (Figure 3E). Differences in bacterial replication cannot account for this effect, as bacteria were quantified shortly after infection, before intracellular replication had begun. Instead, the increase in multi-bacteria infection events in the T110A strain likely results from both the increased fraction of SPI-1(+) cells and enhanced membrane ruffling of epithelial cells. SPI-1(+) cells secrete effector proteins that hijack the host cytoskeleton and induce membrane ruffling, which has been shown to stimulate uptake of SPI-1(+) and nearby SPI-1(−) cells [59]. Together, these results demonstrate that reduced H-NS DNA binding affinity directly affects epithelial cell infection.

### *hns* genotype restricts evolutionary adaptive strategies in SPI-2 inducing environments

The H-NS DBD is highly conserved across enteric bacteria (Figure 1B), indicating strong selection against mutations in this region. Mutations in global regulators such as H-NS may be particularly deleterious as they can have wide-ranging effects – the T110A strain has both fitness defects in SPI-2-inducing and nutrient-limited media (Figures 1A, S1A), as well as altered virulence gene expression (Figures 3B-C). We hypothesized that the *hns* genotype may therefore alter which mutations provide fitness benefits during adaptation to SPI-2-inducing conditions, thereby constraining evolutionary trajectories. In the T110A background, impaired H-NS repression potentially favors mutations that reduce activation of target genes, such as the virulence genes which are misregulated in this strain. Additionally, selection may favor reversion of the T110A mutation or require compensatory mutations elsewhere in the genome. In contrast, H-NS repression is intact in the WT strain, which may favor a distinct set of adaptive mutations.

Comparing evolutionary trajectories between the H-NS WT and T110A strains would reveal whether the starting *hns* genotype constrains the paths available for adaptation. In addition, comparing the magnitude of fitness gains between the strains could indicate whether the WT strain, in which H-NS repression of target genes has been optimized over evolutionary timescales, has limited capacity for fitness gains relative to the T110A strain, in which H-NS function is impaired.

To test this hypothesis, we performed experimental evolution by serially passaging six independent cultures of each strain in SPI-2-inducing media (MgM-MES) for 30 days (Figure 4A) and compared growth metrics, genomic changes, and virulence gene expression. We chose MgM-MES because the T110A strain has a severe fitness defect and altered SPI-2 gene expression in this media. Although WT grows better than T110A in MgM-MES, its growth is still reduced compared to replete media, suggesting there are still opportunities for adaptation. WT and T110A evolved clones displayed distinct growth dynamics (Figures 4A, S4A). At day 5, the WT clones showed no improvement compared to the ancestral WT strain. In contrast, five of six T110A strains had improved fitness and surpassed even the ancestral WT strain, with variable changes in lag time, maximum growth rate, and maximum OD_600_. By day 30, all WT clones had increased fitness relative to the ancestral WT strain with improvements in all growth metrics, although the magnitude of improvement varied between clones. The T110A clones maintained their improved fitness over the ancestral WT strain at day 30. The differences in the timing of fitness improvements (faster adaptation for T110A) suggest that distinct mutational trajectories occur in each *hns* genotype and that *hns* DBD mutants are genetically unstable, similar to Δ*hns* strains [40].

**Figure 4.**
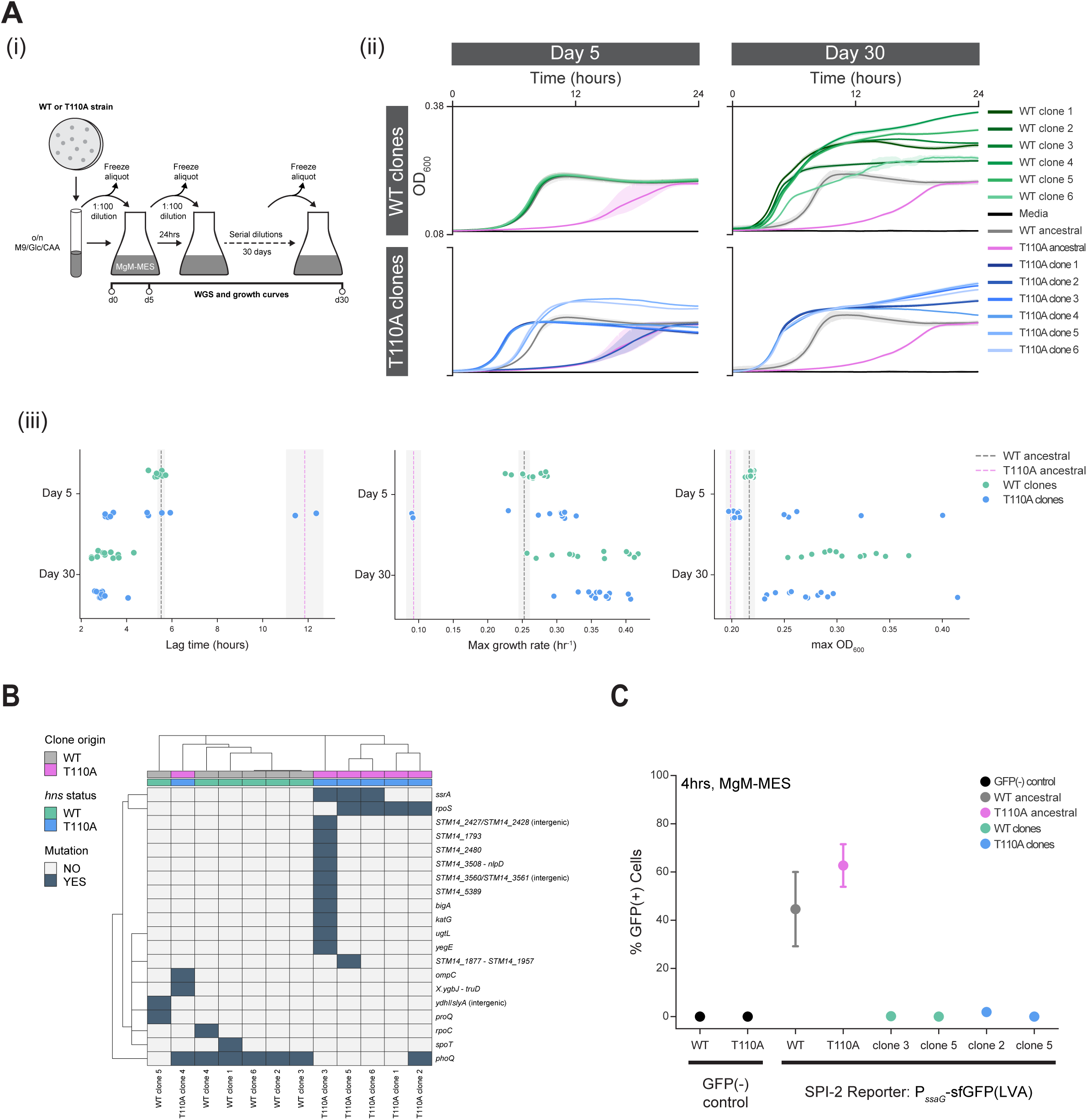
*hns* genotype constrains adaptive mutations that improve fitness in SPI-2 inducing conditions. (A) Experimental evolution reveals fitness gains in both WT and T110A strains. (i) Schematic showing the experimental evolution approach. Six individual colonies from each of the WT and T110A strains were inoculated into M9/Glc/CAA overnight, then subcultured daily in 25 mL SPI-2 inducing media (MgM-MES) for 30 days. Glycerol stocks were made daily. Evolved clones at day 5 (d5) and day 30 (d30) were assessed for growth and genotype using plate reader assays and whole genome sequencing. (ii) Growth dynamics of evolved clones. Most T110A clones (shades of blue) show increased fitness by d5 that stabilizes by d30, while WT clones (shades of green) remain similar to the ancestral WT strain at d5 but show increased fitness by d30. Strains were grown in MgM-MES and OD_600_ was recorded using a plate reader. Growth curves show means of three technical replicates, and shaded regions represent standard deviation. Ancestral strains (WT – gray, T110A – magenta) and media control (black) are shown in each plot for comparison. Representative experiments for d5 and d30 are shown. (iii) Growth metrics extracted from data in (ii) and Figure S4A. Circles represent WT (green) and T110A (blue) evolved clones; dashed lines represent means of the ancestral WT and T110A strains; shading indicates variation between two replicates. (B) WT and T110A evolved clones acquire distinct mutations by d30. Genomic DNA from d30 evolved clones was sequenced and mutations are displayed as a clustered heatmap. WT and T110A clones cluster separately (except T110A clone 4). Colorbars indicate ancestral strain (WT – gray, T110A – magenta) and *hns* genotype at d30 (WT – green, T110A – blue). Dark blue in the heatmap indicates the presence of a mutation in the indicated gene, while light gray indicates no mutation. (C) Evolved clones show negligible SPI-2 induction. SPI-2 induction was assessed in two d30 clones from each background (WT clones 3 and 5; T110A clones 2 and 5) using the SPI-2 GFP reporter plasmid (P_ssaG_-sfGFP(LVA)). Samples were analyzed by flow cytometry after 4 hours in MgM-MES and the percentage of GFP(+) cells compared to that in the ancestral WT and T110A strains. Ancestral strains with GFP(−) control plasmids are shown as a reference for background fluorescence values.

To identify strain-specific adaptive mutations, we performed whole-genome sequencing on all day 30 evolved clones (Figures 4B, S4B). Based on hierarchical clustering of the acquired mutations, WT and T110A clones clustered separately, except for T110A clone 4, which was grouped with the WT clones. Despite the fitness cost of the T110A mutation in MgM-MES, none of the T110A clones reverted to wild type. Instead, all T110A clones acquired compensatory mutations in at least one of *ssrA* (a sensor kinase that activates the SPI-2 master regulator SsrB), *phoQ* (a sensor kinase that activates SPI-2 in response to acidic pH and limited magnesium), or *rpoS* [20,60–64]. The prevalence of *rpoS* mutations is consistent both with H-NS promoting RpoS degradation and the viability of STm Δ*hns* strains often depending on compensatory mutations in *rpoS* [16,18,65]. One T110A clone (clone 3) acquired a mutation in *mutS*, a mismatch repair gene, resulting in numerous additional mutations [66]. In contrast to the T110A clones, all WT clones acquired mutations in genes related to the PhoPQ two-component system. Five clones carried mutations in the sensor kinase *phoQ*, while one carried a mutation in *proQ*, an RNA-binding protein that positively regulates the PhoP regulon [67,68]. While mutations in both WT and T110A clones target the SPI-2 gene regulatory network (GRN), they occur in distinct nodes of the network. Sequencing of three day 5 T110A clones revealed few mutations, consistent with these populations likely being a mixture of cells carrying different mutations that had yet to become fixed in the population (Figure S4B).

To assess whether these mutations in the SPI-2 GRN altered virulence gene expression, we measured SPI-2 reporter expression in two representative day 30 clones from each background (WT clones 3 and 5, T110A clones 2 and 5). After 4 hours in MgM-MES, all four clones showed negligible GFP(+) cells by flow cytometry (Figures 4C, S4C). In three of the clones, the small fraction of GFP(+) cells had intensity measurements similar to those of rare cells in the GFP(−) control strain labeled as GFP(+), confirming that these cells are unlikely to be SPI-2 responders (Figure S4C). Only T110A clone 2 had GFP(+) cells of similar intensity to the ancestral strains, indicating that this clone retained SPI-2 activity, albeit in a much decreased fraction of cells.

Overall, the genomic changes observed in the evolved clones confirm that distinct adaptive mutations occur in each *hns* genotype, suggesting that T110A clones must compensate for the broader effects of H-NS target gene misregulation. Together, these results reveal how evolutionary constraints on a highly conserved global regulator impact pathogen adaptation to intracellular-like environments.

## DISCUSSION

Here, we characterized an STm strain with a mutation (T110A) in the H-NS DBD to assess the contribution of H-NS-mediated repression to virulence gene expression heterogeneity and pathogen adaptation to intracellular-like environments. Biochemical characterization confirmed the T110A mutation reduced H-NS DNA-binding affinity *in vitro*. Using SPI-1 and SPI-2 transcriptional reporters, we found that this reduced H-NS DNA-binding affinity increased the fraction of responding cells while preserving bimodal expression of the virulence reporters. The increased fraction of SPI-1(+) cells in the T110A strain had functional consequences, resulting in increased infection of epithelial cells *in vitro*. Finally, using experimental evolution we demonstrate that *hns* genotype constrains adaptive mutations to intracellular-like conditions, with distinct mutations associated with fitness gains in each strain. Together, our findings establish that the global repressor H-NS contributes to virulence gene expression heterogeneity by modifying virulence activation thresholds in individual bacteria, while strong purifying selection on the H-NS DBD restricts the mutational paths available during adaptation to intracellular-like environments.

Optimal H-NS-mediated repression of virulence genes requires both tight silencing in non-inducing environments and the rapid relief of this silencing upon exposure to inducing conditions. The critical role of H-NS in silencing virulence gene expression in non-inducing environments has been well-established using Δ*hns* strains, while elegant biochemical studies have characterized the role of response regulators in displacing H-NS from virulence gene promoters to enable expression [16,18,23,24,40]. However, whether H-NS-mediated repression modulates heterogeneity in virulence gene expression in inducing conditions has remained unclear for several reasons. First, evaluating the role of a global regulator like H-NS in virulence heterogeneity is challenging because Δ*hns* strains cause widespread gene dysregulation and severe growth defects even in rich media, rapidly selecting for compensatory mutations that confound interpretation of the contribution of H-NS to any phenotype [16,18,40]. Second, in Δ*hns* strains, several STm virulence regulons are broadly expressed in non-inducing environments, precluding analysis of how relief of H-NS repression impacts virulence heterogeneity in inducing conditions [16,18]. The H-NS T110A mutant overcomes both challenges and, as such, is ideally suited to evaluate how impaired H-NS function impacts virulence gene expression heterogeneity. In non-inducing conditions, the T110A strain grows normally, limiting selective pressure for compensatory mutations, and maintains virulence gene repression, enabling the role of H-NS in virulence gene activation to be directly evaluated.

The preservation of bimodality in both SPI-1 and SPI-2 responses in the T110A strain demonstrates that H-NS itself does not determine whether heterogeneity occurs, but instead it modifies features of the virulence response at the single-cell level. For both SPI-1 and SPI-2, the fraction of responders is the feature most affected by reduced H-NS DNA-binding affinity (Figure 3). While the activation of response regulators sets an activation threshold that individual bacteria must exceed to express virulence genes, our results indicate that H-NS repression can adjust this threshold, at least in a subset of cells. Two models to explain the increase in the fraction of virulence responders in the T110A strain are: 1) response regulator levels vary continuously across the population, and in certain bacteria these levels are insufficient to displace H-NS WT but sufficient to displace the reduced affinity H-NS T110A, or 2) H-NS repression of virulence genes varies between individual bacteria, such that weakening H-NS repression shifts only a subset of cells past the activation threshold. These models make distinct, testable predictions, although testing them will require technically challenging measurements such as determining the relationship between levels of response regulators and virulence gene expression in individual bacteria or quantifying variation in H-NS deposition at virulence gene promoters in single bacteria.

STm has evolved to navigate fluctuating host environments and adjust virulence gene expression accordingly [3,4]. As such, virulence gene expression is a dynamic process in which the coordinated interplay between H-NS and response regulators is ongoing as the pathogen cycles between modes of virulence gene activation and repression based on environmental cues. Examining virulence responses in single cells over time has revealed critical features of the SPI-2 response, including switch-like activation and heterogeneity in response timing [33]. An important future extension of our findings will be to examine virulence gene expression over time in individual bacteria with either intact or impaired H-NS repression, both *in vitro* and in host cells. The H-NS T110A mutation may have consequences that extend beyond the fraction of virulence responders. For instance, the WT and T110A strains could differ in the trajectory of the virulence response, the stability of expression across generations, or the reversibility of the response as cells transition to new environments, all processes that involve either the relief or reestablishment of H-NS repression. Such studies will be critical to gain mechanistic insight into how H-NS repression of virulence genes has been optimized over evolutionary time.

H-NS repression plays a critical role in the maintenance of virulence genes and thus enables *Salmonella* colonization of diverse host environmental niches [16,18,29,36,40,69]. However, whether H-NS itself, as a result of its highly conserved DBD, constrains mutations during adaptation to new host environments was unclear. Combining experimental evolution with the T110A strain enabled us to demonstrate that evolutionary trajectories in SPI-2 inducing conditions differed in several key ways depending on the starting *hns* regulatory state. While both WT and T110A strains converged on disrupting the SPI-2 regulatory network, mutations arose in different nodes in the network depending on the starting *hns* genotype. T110A clones likely selected for mutations that simultaneously improve fitness and compensate for baseline gene expression dysregulation, whereas WT clones, lacking this dysregulation could select for a distinct set of mutations. Revertants restoring WT H-NS were never observed, suggesting that if compensatory mutations restore fitness, reversion is unlikely, consistent with simulations of reversion frequency in bacteria [70]. Differences in the dynamics of adaptation revealed that the context-specific growth defects of the T110A strain rapidly select for compensatory mutations in SPI-2 inducing conditions, consistent with strong purifying selection on the H-NS DBD (Figures 1, 4). Finally, in both *hns* genotypes, adaptation came at a cost, the loss of SPI-2 inducing capacity. This indicates a tradeoff between fitness and maintenance of intact virulence gene regulation and suggests that SPI-2 expression incurs a fitness cost, at least under prolonged exposure to inducing conditions. Although technically challenging to implement, future experimental evolution approaches should incorporate periodic selection for functional SPI-2 to reveal whether bacterial populations can simultaneously optimize growth and maintain inducible virulence gene regulation, or whether this is a fundamental constraint that limits pathogen adaptation.

In summary, we find that evolutionary constraints on the global regulator H-NS restrict both heterogeneity in virulence gene expression and pathogen adaptation to intracellular-like environments. Our findings demonstrate that optimization of H-NS repression extends beyond tight silencing of virulence genes in non-inducing environments to include rapid and heterogeneous activation upon exposure to intracellular-like conditions, and that H-NS DBD mutants will be a valuable resource to further dissect this process.

## ACKNOWLEDGEMENTS

We thank members of the Lane lab for discussions and valuable feedback on the manuscript. We thank members of the Blythe, Brickner, Horvath, Lackner, Mondragón, and Weiss labs for advice, reagents, and technical assistance with protein purification and EMSAs. We thank the Robert H. Lurie Comprehensive Cancer Center Flow Cytometry Core Facility of Northwestern University and the Single Cell Genomics Core for flow cytometry training and equipment. The Lurie Cancer Center is supported in part by an NCI Cancer Center Support Grant #P30 CA060553. We thank the Northwestern Center for Synthetic Biology Biofoundry for training and use of the AKTA Avant 25 for SEC. UCSF ChimeraX was developed by the Resource for Biocomputing, Visualization, and Informatics at the University of California, San Francisco, with support from National Institutes of Health R01-GM129325 and the Office of Cyber Infrastructure and Computational Biology, National Institute of Allergy and Infectious Diseases. We acknowledge our funding sources: NIH Cellular and Molecular Basis of Disease Training Program award to MRS (NIH T32 GM008061) and a Northwestern Weinberg College of Arts and Sciences Baker Faculty Research Program award to K.L.

## AUTHOR CONTRIBUTIONS

Conceptualization: MS, KL. Data curation: LM, DK, KL. Formal analysis: LM, DK, MS, MB, JZ, KL. Funding Acquisition: MS, KL. Investigation: LM, DK, MS, MB, JZ, KL. Methodology: LM, MS, KL. Project Administration: KL. Resources: MS, KL. Supervision: KL. Validation: KL. Visualization: LM, DK, MS, KL. Writing – Original Draft Preparation: LM, MS, KL.

## METHODS

Essential reagents and catalog numbers are listed in Table S3.

### Salmonella strains

The *Salmonella enterica* serovar Typhimurium 14028 strain used as wild type (WT) was a gift from the laboratory of Michael McClelland. The *hns* mutant strain was isolated from a preceptrol freeze-dried stock of *Salmonella enterica* serovar Typhimurium 14028 obtained from ATCC (ATCC 14028). The strain was rehydrated and streaked on nutrient agar according to ATCC protocols. Colonies from the initial streak varied in morphology, and several appeared mucoid. Individual colonies were restreaked onto LB agar plates and glycerol stocks were prepared. One glycerol stock was used for all experiments and is referred to as the *hns* mutant or T110A strain throughout.

### *Salmonella* Cell Culture

Bacterial strains were streaked from glycerol stocks onto LB agar containing appropriate selective antibiotics and incubated overnight at 37°C. Plates were used for experiments within one week and culture growth kept to a minimum to avoid issues of genomic instability associated with *hns* mutant strains [40]. Cultures were grown at 37°C with shaking (250 rpm). Growth media compositions: LB-Lennox (LB): NaCl 5g/L; LB-Miller (LB High Salt): NaCl 10g/L (SPI-1 inducing media); M9/Glc/CAA: 1x M9 salts, 0.1mM CaCl_2_, 2mM MgSO_4_, 0.4% glucose, 0.2% casamino acids; M9/Glc: 1x M9 salts, 0.1mM CaCl_2_, 2mM MgSO_4_, 0.4% glucose; MgM-MES (SPI-2 inducing media): 5mM KCl, 7.5mM (NH_4_)_2_SO_4_, 0.5mM K_2_SO_4_, 1mM KH_2_PO_4_, 170mM 2-(N-morpholino)ethanesulphonic acid hydrate (pH adjusted to 5.0 with NaOH), 0.1% casamino acids, 38mM glycerol, 8µM MgCl_2_ [71,72]. Where indicated, glycerol was replaced with 0.4% glucose.

### Plasmids, Cloning, and *Salmonella* Strain Generation

All plasmids and strains generated in this study are listed in Table S4. Plasmids were assembled using either Golden Gate Assembly with the EcoFlex library or Gibson assembly [73]. *E. coli* TG1 cells were used for all cloning steps. Plasmids were verified by Sanger sequencing or whole plasmid sequencing.

*SPI-1 reporter plasmid (P*_invF_*-sfGFP):* The *invF* promoter region (340 bp upstream of the start codon plus 179 bp of the coding region) was previously described [53] and was amplified from STm genomic DNA using PrimeSTAR Max DNA polymerase and cloned into a Golden Gate Level 0 promoter plasmid using Gibson assembly. A Level 1 single transcriptional unit (TU) plasmid containing the promoter, ribosome binding site, open reading frame, and terminator was assembled by Golden Gate Cloning. This plasmid was combined with a second TU expressing a constitutive mRuby2 to generate the final Level 2 plasmid.

*SPI-2 reporter plasmid (PssaG-sfGFP-LVA) and constitutive mRuby2 plasmid (P*_SJM901_*-mRuby2):* These plasmids were generated using the same strategy and have been previously described [33].

*Strain construction:* Electrocompetent cells were prepared using the mannitol-glycerol preparation method [74]. Reporter plasmids were used to transform bacteria, and transformants were selected on LB agar plates containing the appropriate antibiotic (see Table S4.2). WT strains containing the SPI-2 reporter plasmid or the constitutive mRuby2 plasmid were previously generated [33]. Glycerol stocks were prepared by mixing 700µl overnight culture (grown in LB with appropriate antibiotics) with 300µl of 50% w/v glycerol, vortexing, and storing at –80°C.

### Mammalian Cell Culture

HeLa cells from ATCC were maintained in DMEM supplemented with 10% FBS, 1x Penicillin/Streptomycin, and 2mM GlutaMAX at 37°C, 5% CO_2_. Cell lines were not authenticated.

### Generation of Stable Cell Lines

Lentivirus was produced in Lenti-X 293T cells using packaging plasmids psPAX2 (Addgene #12260) and pMD2.g (Addgene #12259). HeLa cells were transduced with lentivirus expressing H2B-miRFP670 and selected with hygromycin.

### Plate Reader Growth Assays and Analysis

Single colonies were inoculated into the appropriate pre-culture medium (M9/Glc/CAA for M9/Glc/CAA, M9/Glc, and MgM-MES plate reader samples and LB-Lennox for LB-Lennox and LB-Miller plate reader samples) and grown shaking at 37°C for 14-16 hours. Overnight cultures were backdiluted to OD_600_ of 0.06 in 3 mL of the same media and grown shaking for 3 hours. Cells were pelleted, washed twice in the plate reader media, and diluted to OD_600_ of 0.002. Three technical replicates (150µl each) were added to a glass-bottom 96-well black plate. Plates were sealed with breathable film, and a needle was used to pierce each well to ensure aeration. Following a 30-minute equilibration in the plate reader (Tecan Spark) at 37°C, OD_600_ was measured at 5-minute intervals for 24 hours. Growth curve metrics were extracted using the Omniplate software package [75]. Plots were generated using Seaborn and Matplotlib in Python [76,77]. Statistical analyses were performed in Python (scipy and statsmodels) using unpaired t-tests. Asterisks denoting significance level are indicated in the Figure legends.

### Whole Genome Sequencing (WGS) and Analysis

Strains were streaked onto LB agar plates and incubated overnight at 37°C. Single colonies were inoculated into 3 mL LB and grown shaking overnight at 37°C. Genomic DNA was isolated from 0.5-1 mL of culture using the Wizard(R) Genomic DNA purification kit (Promega) and eluted in Tris pH 8.5. DNA samples were processed for Illumina whole-genome sequencing (200 Mbp sequencing) by SeqCenter (https://www.seqcenter.com/).

Sequencing reads were analyzed using *breseq* [78]. FASTQ files were aligned to the STm 14028s reference genome, including the chromosome (GenBank: CP001363.1) and plasmid (GenBank: CP001362.1) sequence files. All samples were compared to the STm WT strain used in the lab. The mutation heatmap (Figure 4B) was generated in R using the pheatmap package with binary distance clustering (clustering_distance_cols = ‘binary’) and complete linkage (clustering_method = ‘complete’). A complete list of genetic changes for all WGS samples is reported in Table S1.

### Protein Sequence Alignment

Protein sequences of H-NS homologs from Gram-negative enteric bacteria were retrieved from UniProt. VicH, an H-NS-like protein from *Vibrio cholerae*, was also included [79]. Sequences were aligned using Clustal Omega with output order set to match input order [80]. Alignments were visualized using Jalview with color set to percent identity and conservation scores displayed [81].

### Protein Structure Analysis

Structural analysis was performed in UCSF ChimeraX using the NMR-resolved structure of the STm H-NS DNA-binding domain (PDB ID: 2L93) [15,82]. The T110A mutant model was generated by substituting alanine for threonine at position 110 in the wild-type structure. Wild type and T110A models were compared to evaluate local structural effects of the substitution around the QGR motif. Hydrogen bonds were displayed, and van der Waals interactions were identified by displaying contacts within 3.5 Å.

### Protein Expression and Purification

Expression plasmids for H-NS WT and T110A were generated in the pET-28a backbone with a C-terminal PreScission protease (PPX) cleavage site and 6xHis tag (Figure S2A). Expression plasmids were used to transform competent *E. coli* BL21 (DE3) cells, and transformants were selected on LB agar with kanamycin (50 µg/mL). Cultures were grown in LB-Lennox with kanamycin to OD_600_ of 0.5, induced with 0.5 M isopropyl-beta-D-thiogalactopyranoside (IPTG), and grown for 2 hours at 37°C with shaking (250 rpm). Cells were pelleted (14,000 rpm, 30 minutes, 4°C), resuspended in lysis buffer (20mM Tris-HCl pH 8, 150mM NaCl, 5% v/v glycerol, 2mM 1,4-dithiothreitol (DTT), 1mM phenylmethylsulfonyl fluoride (PMSF), 1x EDTA-free protease inhibitor) and sonicated on ice (Branson, output control of 2.5%, 20% duty cycle, 5-second pulses with 45-second rest intervals on ice). Lysates were clarified by centrifugation (14,000 rpm, 30 minutes, 4°C). All subsequent precipitation steps were performed at 4°C. DNA was precipitated out of clarified lysates by adding polyethyleneimine (PEI; avg. MW 60 K) slow and drop-wise: 0.4% PEI in 150mM NaCl for H-NS WT, and 0.35% PEI in 1M NaCl for H-NS T110A. Samples were incubated for 15 minutes with gentle stirring and then centrifuged (11,000*xg*, 15 minutes, 4°C). The supernatant was gently decanted into a beaker, and H-NS was salted out by slow addition of powdered ammonium sulfate to 37% saturation. The solution was stirred gently overnight, and precipitated H-NS was collected by centrifugation (15,000 rpm, 40 minutes, 4°C). The H-NS precipitate was carefully resuspended in 10 mL of binding buffer (20mM Tris-HCl pH 8, 300mM NaCl, 20mM imidazole, 5mM β-mercaptoethanol). Samples were applied to pre-equilibrated Ni-NTA agarose resin (Invitrogen) and incubated on ice with gentle rocking for 1 hour. Resin was pelleted (800*xg,* 1 minute, 4°C), and the flow-through was discarded. To reduce non-specific binding and disrupt remaining protein-DNA interactions, the resin was washed with wash buffer (20mM Tris-HCl pH 8, 600mM NaCl, 50mM imidazole, 5mM β-mercaptoethanol) for 1 hour on ice with gentle rocking, then pelleted (800*xg*, 1 minute, 4°C). Wash buffer was decanted, and cleavage buffer was added (20mM Tris-HCl pH 8, 300mM NaCl, 5mM β-mercaptoethanol) along with PreScission Protease (1µl per nmol H-NS). Samples were incubated on ice with gentle rocking for 4 hours. Resin was pelleted (800*xg*, 1 minute, 4°C) and the H-NS-containing flow-through was collected. Samples were concentrated using a 3kDa MWCO cutoff Ultra Centrifugal Filter (Amicon) and subjected to size-exclusion chromatography (SEC) on a Superdex 75 Increase 10/300 GL column (Cytiva) equilibrated in SEC buffer (20 mM Tris-HCl pH 8.0, 300 mM NaCl, 5% glycerol, 5 mM β-mercaptoethanol) and run at a flow rate of 0.8 mL/min on an ÄKTA Avant 25 system (GE Healthcare) at Northwestern University’s Biofoundry Core. Peak fractions were analyzed by Coomassie-stained SDS–PAGE and UV–Vis spectrophotometry. Protein purity was quantified from the Coomassie-stained gel using ImageJ [83] and determined to be ∼90% for H-NS WT and 99% for H-NS T110A. Absorbances at 260 nm and 280 nm were used to verify protein concentration and absence of nucleic acids. A_260_/A_280_ ratios for all samples were approximately 0.6, consistent with minimal nucleic acid contamination. Samples were flash-frozen in liquid nitrogen and stored at –80°C.

### Electrophoretic Mobility Shift Assays (EMSAs) and Analysis

A region of the *ssrAB* promoter region was amplified from STm genomic DNA using PrimeSTAR DNA Polymerase (Takara) with primers previously described (Figure S2C) [45]. PCR products were purified (Zymo Research) and verified by gel electrophoresis and Sanger sequencing (Azenta). WT and T110A H-NS proteins were exchanged into EMSA binding buffer (20mM Tris-HCl pH 7.4, 4mM MgCl_2_, 100mM NaCl, 1mM DTT) using 3kDa MWCO cutoff Ultra Centrifugal Filters (Amicon). Binding reactions were assembled in 0.2 mL reaction tubes. Water, then EMSA Binding Buffer was added to the desired volumes and mixed. Protein was added (0, 200, 350, 450, 550, 600, 650, 750, 850, 950nM, 1, 1.5, 2, 3µM) and gently mixed. In 30-second increments, 25ng DNA probe per reaction tube was added. Samples were incubated at room temperature for 30 minutes. Loading dye (40% glycerol, 0.25% bromophenol blue) was added, and samples were loaded onto a 5% native Mini-PROTEAN TBE gel (Bio-Rad) in 30-second increments to maintain consistent timing across reactions. Gels were run at 100V in 0.5x TBE buffer for 30 minutes, stained with Sybr Safe (Invitrogen), and washed twice with 150 mL Milli-Q H_2_O. Gels were imaged on an Azure 600 (Azure Biosystems Inc.) with a 10-second exposure time.

Gels were quantified using ImageJ. Background subtraction was performed by measuring the median intensity of four background regions and calculating the mean. Bound and unbound DNA regions were defined for each lane on a gel using the ROI manager tool. The fraction bound was calculated for each lane as: fraction bound = (bound) / (bound + unbound), and a min-max normalization was performed. All calculations and non-linear regression fitting to the Hill equation were performed using SciPy and Python.

### SPI-1 and SPI-2 Induction Assays

*SPI-1 induction:* Single colonies were inoculated into 3 mL LB-Lennox with appropriate antibiotics and grown shaking at 37°C for 14-16 hours. Overnight cultures (BD1) were back diluted to OD_600_ of 0.06 in fresh LB-Lennox and grown shaking for 2.5 hours (BD2). Cultures were then back diluted to an OD_600_ of 0.06 into 5 mL LB-Miller (SPI-1 inducing) or LB-Lennox (control). At the indicated fixation timepoints, 600µl of culture was mixed with 200µl of 16% paraformaldehyde by inversion and incubated at room temperature for 30 minutes. Fixed samples were washed three times with 1 mL PBS before storing in 1 mL PBS at 4°C. Samples were analyzed by flow cytometry within one week of fixation.

*SPI-2 induction:* Single colonies were inoculated into 3 mL M9/Glc/CAA with appropriate antibiotics and grown shaking at 37°C for 14-16 hours. Overnight cultures were back diluted 1:100 into 3 mL fresh M9/Glc/CAA and grown shaking for 4 hours. Cultures were then back diluted to an OD_600_ of 0.02 in 5 mL of MgM-MES containing either glycerol or glucose. Samples were fixed and stored as described for SPI-1 induction.

### Flow Cytometry and Analysis

Fixed samples were diluted in PBS to achieve approximately 10,000 events per second or below. Samples were analyzed on a BD LSRFortessa SORP Cell Analyzer with HTS (6-laser 18-parameter). At least 50,000 mRuby2 positive events were captured per sample.

Data were analyzed using FlowJo v10.8.2. Forward scatter (FSC) and side scatter (SSC) gates were drawn to capture the majority of events. All reporter strains constitutively express mRuby2; therefore, mRuby2 positive gates were set using negative control samples (cells without the mRuby2 plasmid) such that less than 0.05% of the negative control events were classified as mRuby2 positive. mRuby2 positive events were downsampled to 50,000 using the FlowJo Downsample plugin (V3.3.1) for all subsequent analyses. GFP-positive gates were set using a GFP-negative control sample (strain with a promoterless GFP control plasmid). Statistics, including mean fluorescence intensity (MFI) and percentage of GFP(−) and GFP(+) cells, were exported from FlowJo. Plots were generated using Seaborn and Matplotlib in Python. Statistical analyses were performed in Python (scipy and statsmodels). Comparisons of percent GFP(+) cells between strains were analyzed using a linear mixed-effects model with strain, time and media as factors and accounting for repeated measures from the same culture tube. Post-hoc pairwise comparisons between WT and T110A strains within each media-time combination (SPI-2 data: 4 total (2 media, 2 timepoints), SPI-1 data: 6 total (2 media, 3 timepoints)) were performed using unpaired t-tests with p values corrected for multiple comparisons using the Holm-Sidak method. Consistency in trends across replicates was assessed using the sign test. Asterisks denoting significance level are indicated in the Figure legends.

### Fluorescence Microscopy

HeLa cells were plated at 6,000 cells/well on fibronectin-coated (10µg/ml) 96-well glass bottom plates one day prior to infection. Bacterial cultures were prepared as described in the SPI-1 induction assays: Single colonies were inoculated into 3mL LB-Lennox with appropriate antibiotics and grown shaking at 37°C for 14-16 hours. Overnight cultures were back diluted to an OD_600_ of 0.06 in fresh LB-Lennox and grown shaking for 2.5 hours, then back diluted to an OD_600_ of 0.06 in 5 mL LB-Miller (SPI-1 inducing) or LB-Lennox (control) and grown shaking for 4 hours. One hour prior to infection, HeLa cells were washed three times with imaging media (1x FluoroBrite DMEM, 1% FBS, 2 mM GlutaMAX) to remove residual Penicillin/Streptomycin and 100µl of imaging media was added to each well. Bacterial cultures were pelleted, resuspended in PBS, and diluted to the indicated multiplicity of infection (MOI). 5µl of bacteria was added to each well and the plate was incubated at 37°C for 30 minutes. HeLa cells were washed three times with imaging media to remove extracellular bacteria, and 200µl of imaging media containing 10 µg/mL gentamicin was added to each well. Plates were sealed with an AeraSeal film and imaged. Imaging was performed with a Nikon Ti2E fluorescence microscope equipped with a Prime BSI sCMOS camera, a SPECTRA III light engine, temperature (37°C) and environmental (5% CO_2_) control. Images were acquired using a 20x/0.95 numerical aperture objective with 2×2 binning. Acquisition was controlled by Nikon Elements software.

### Image Analysis

Nuclei labeled with H2B-miRFP670 were segmented using Cellpose3 and a custom trained model [84,85]. Images with sparse nuclei were poorly segmented due to Cellpose’s standard image normalization and were manually identified and excluded from analysis. All segmented images were manually inspected, and missegmented nuclei were removed. Bacteria labeled with mRuby2 were preprocessed to subtract background and reduce noise using 1) N4 bias field correction algorithm, 2) wavelet-based background subtraction, and 3) anisotropic diffusion [86,87]. Nuclei labels were filtered to remove objects <266 pixels in area (1st percentile of all nuclei). To minimize detection of out-of-focus extracellular bacteria, images were further processed using background subtraction (napari-segment-blobs-and-things-with-membranes: nsbatwm) followed by Laplacian of Gaussian filtering (napari-pyclesperanto-assistant) [88]. Bacteria were segmented using Voronoi-Otsu labeling (pyclesperanto-prototype). A minimum threshold for bacteria object intensity (2500 AU) was determined from uninfected control wells. Bacterial objects were assigned to the nearest nucleus using a Euclidean distance threshold of 60 pixels. Features of nuclei and bacterial objects were extracted using scikit-image regionprops and stored in xarray format. Plots were generated using Seaborn and Matplotlib in Python. Statistical analyses were performed in Python (scipy and statsmodels) using two-way ANOVA with strain and MOI as factors.

### Experimental Evolution

Glycerol stocks of WT and T110A strains were streaked on LB agar and incubated overnight at 37°C. Six independent colonies from each strain were inoculated into 3 mL M9/Glc/CAA and grown shaking overnight at 37°C. The following day, 1 mL of each culture was pelleted (6000rpm, 2 minutes) and washed twice in 1 mL MgM-MES to remove residual M9/Glc/CAA. Washed cells were diluted 1:100 (150µl in 15 mL MgM-MES in 125 mL flasks) and grown shaking at 37°C for approximately 24 hours. For 30 days, 150µl of each culture was transferred daily into 15 mL fresh MgM-MES (1:100) without washing. Glycerol stocks were prepared daily to facilitate subsequent growth assays and whole-genome sequencing.

### Statistical Analysis

*Growth curves:* Pairwise comparisons used two-sided, unpaired t-tests and p values were adjusted for multiple comparisons using the Benjamini-Hochberg FDR method. *Flow cytometry:* Comparisons of % GFP(+) cells used a linear mixed-effects model with post-hoc pairwise comparisons performed with unpaired t-tests and Holm-Sidak correction for multiple comparisons. Consistency in trends across replicates was assessed using sign tests. *HeLa cell infection:* Comparisons of infected cell frequency and number of multi-bacteria infection events were performed using two-way ANOVA with strain and MOI as factors. Statistical analyses were performed in Python (scipy and statsmodels). Alpha set to 0.05. All statistical analyses are reported in Table S2.

---

**Figure S1.**
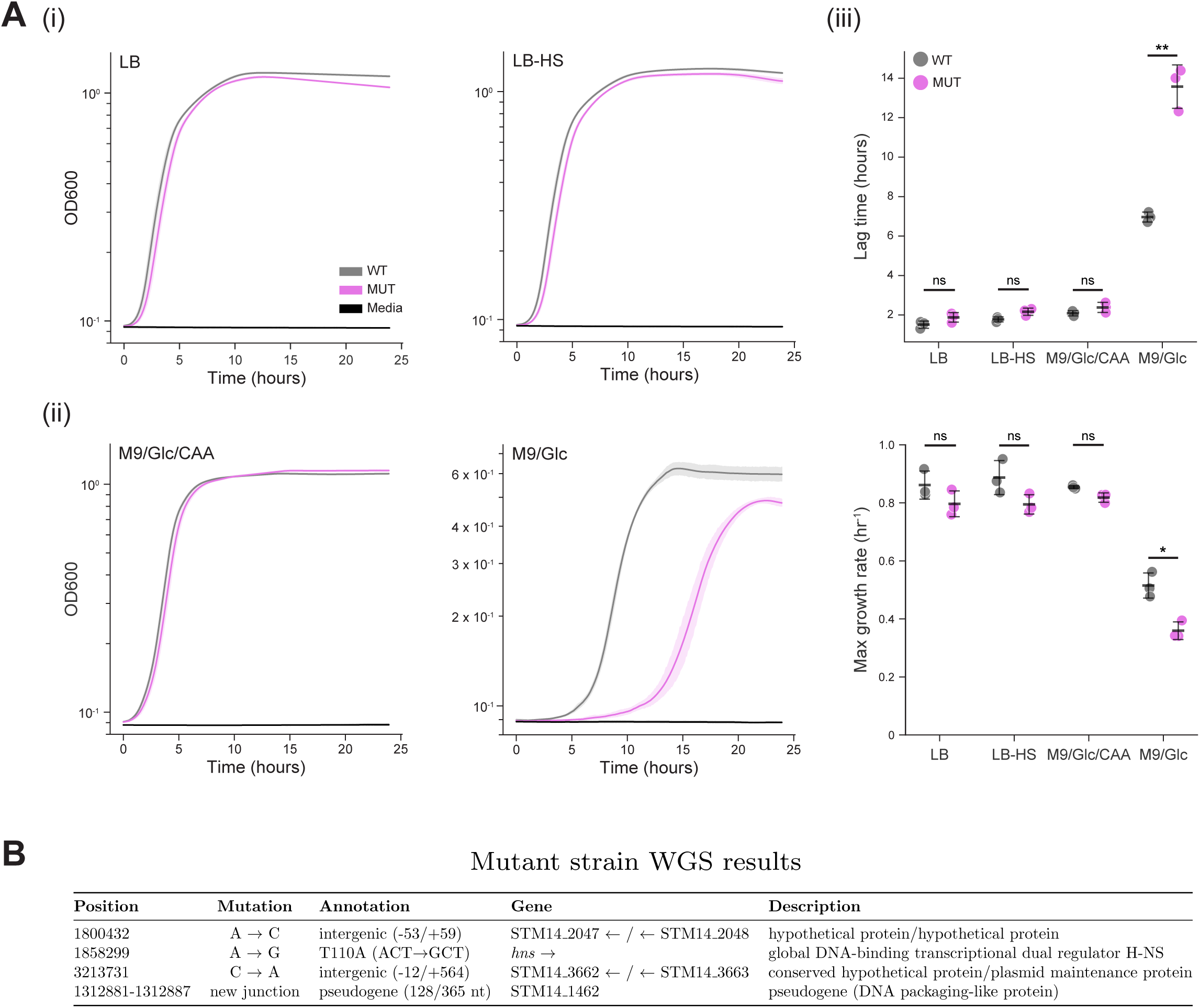
Growth metrics and whole genome sequencing of the mutant strain. (A) Growth curve analysis in additional media conditions. Plate reader growth curves of WT (gray) and MUT (magenta) strains in (i) non-SPI-1-inducing (LB) and SPI-1-inducing (LB-High Salt [LB-HS]) media, and (ii) replete media (M9/Glc/CAA) and nutrient-limited media (M9/Glc). Growth curves show averages of three independent experiments, each with three technical replicates; shaded regions represent standard deviation. Media control is shown in black. Y-axes are scaled to the growth condition. (iii) Growth metrics extracted from the data in (i) and (ii). Individual datapoints represent independent experiments. Asterisks denote significance as determined by two-sided unpaired t-test with Benjamini-Hochberg FDR correction as follows: * p<0.05, ** p<0.01; ns, not significant. (B) Whole genome sequencing identified *hns* a328g (T110A) as the only coding region difference between the WT and MUT strains.

**Figure S2.**
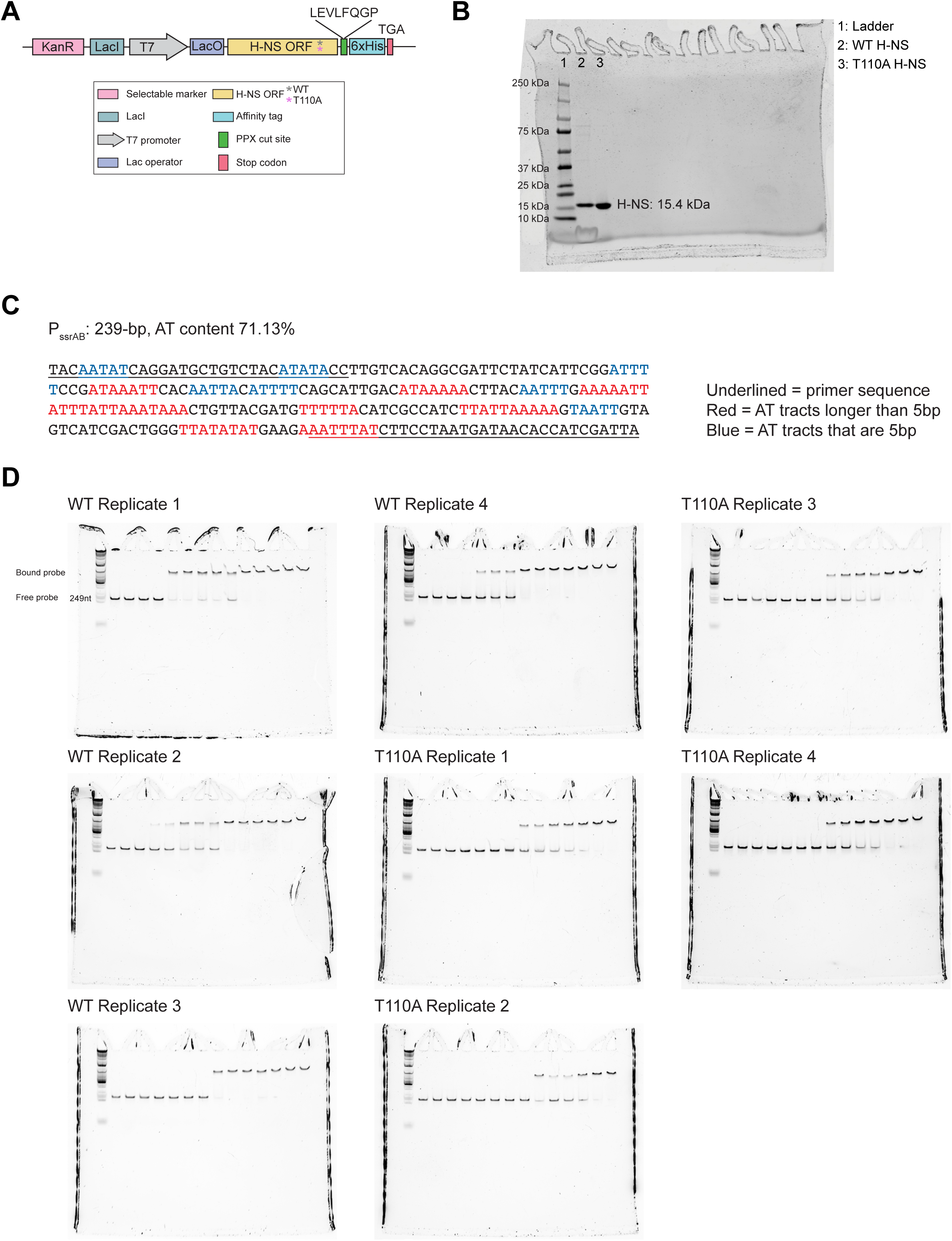
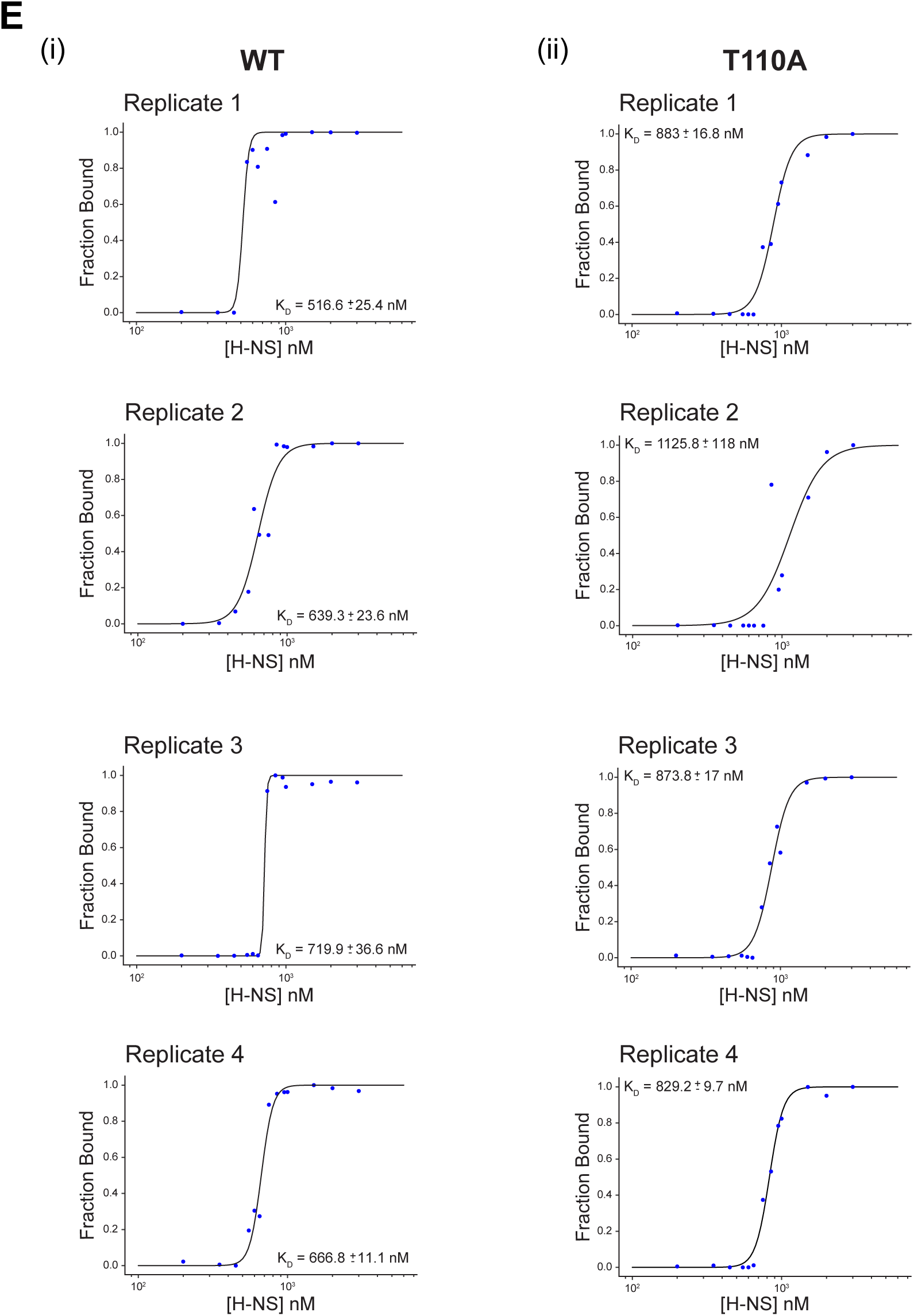
Biochemical characterization of H-NS T110A. (A) Schematic of the plasmid construct used for H-NS protein expression. (B) Full SDS-PAGE gel image corresponding to Figure 2B. (C) Sequence and features of the *ssrAB* promoter region used in EMSAs. Underlined sequences indicate PCR primers used to amplify the region from STm genomic DNA. Colored text highlights AT-rich tracts of potential H-NS binding sites (red: longer tracts, blue: shorter tracts). (D) Complete gel images from four independent EMSA experiments for H-NS WT and T110A proteins. Data from replicate 4 is shown in Figure 2C. (E) Quantification and curve fitting of EMSA data from panel D. Blue circles represent experimental data. Lines represent fits to the Hill equation. K_D_ values are reported as best fit ± SE (standard error).

**Figure S3.**
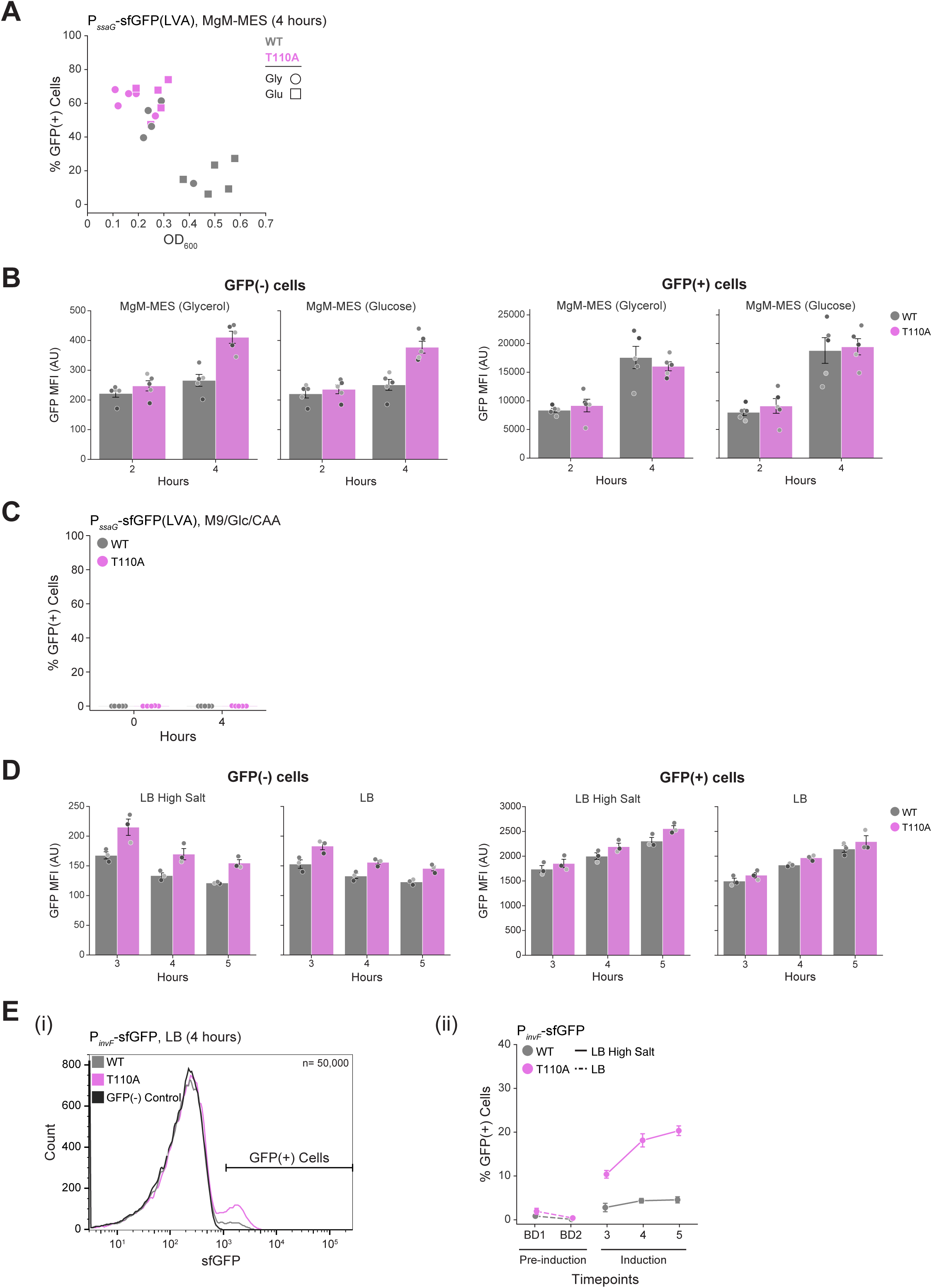
SPI-2 and SPI-1 reporter expression in H-NS T110A strain. (A) SPI-2 reporter (P_ssaG_-sfGFP(LVA)) expression vs growth. Scatter plot shows % GFP(+) cells versus OD_600_ for WT and T110A samples at 4 hours (corresponding to data in Figure 3B) in MgM-MES with glycerol (circles) or glucose (squares) as the carbon source. (B) Mean fluorescence intensity (MFI) of GFP(−) and GFP(+) subpopulations for SPI-2 reporter data shown in Figure 3B. (C) SPI-2 reporter expression under non-inducing conditions. Flow cytometry data was analyzed as in Figure 3B. n=50,000 cells/sample, 5 independent experiments. (D) MFI of GFP(−) and GFP(+) subpopulations for SPI-1 reporter data shown in Figure 3C. (E) SPI-1 reporter (P_invF_-sfGFP) expression in the T110A strain in non-inducing media. (i) Representative histogram of WT and T110A strains containing the SPI-1 reporter plasmid grown in LB for 4 hours. n=50,000 cells. Data corresponds to Figure 3C. (ii) The frequency of SPI-1 reporter GFP(+) cells before induction was assessed by measuring % GFP(+) cells after the first (BD1) and second (BD2) back dilution in LB and comparing to samples grown in SPI-1-inducing media (LB High Salt) for 3, 4, and 5 hours. SPI-1 reporter expression is negligible before induction in LB High Salt. WT strain shown as gray and T110A as magenta across all panels.

**Figure S4.**
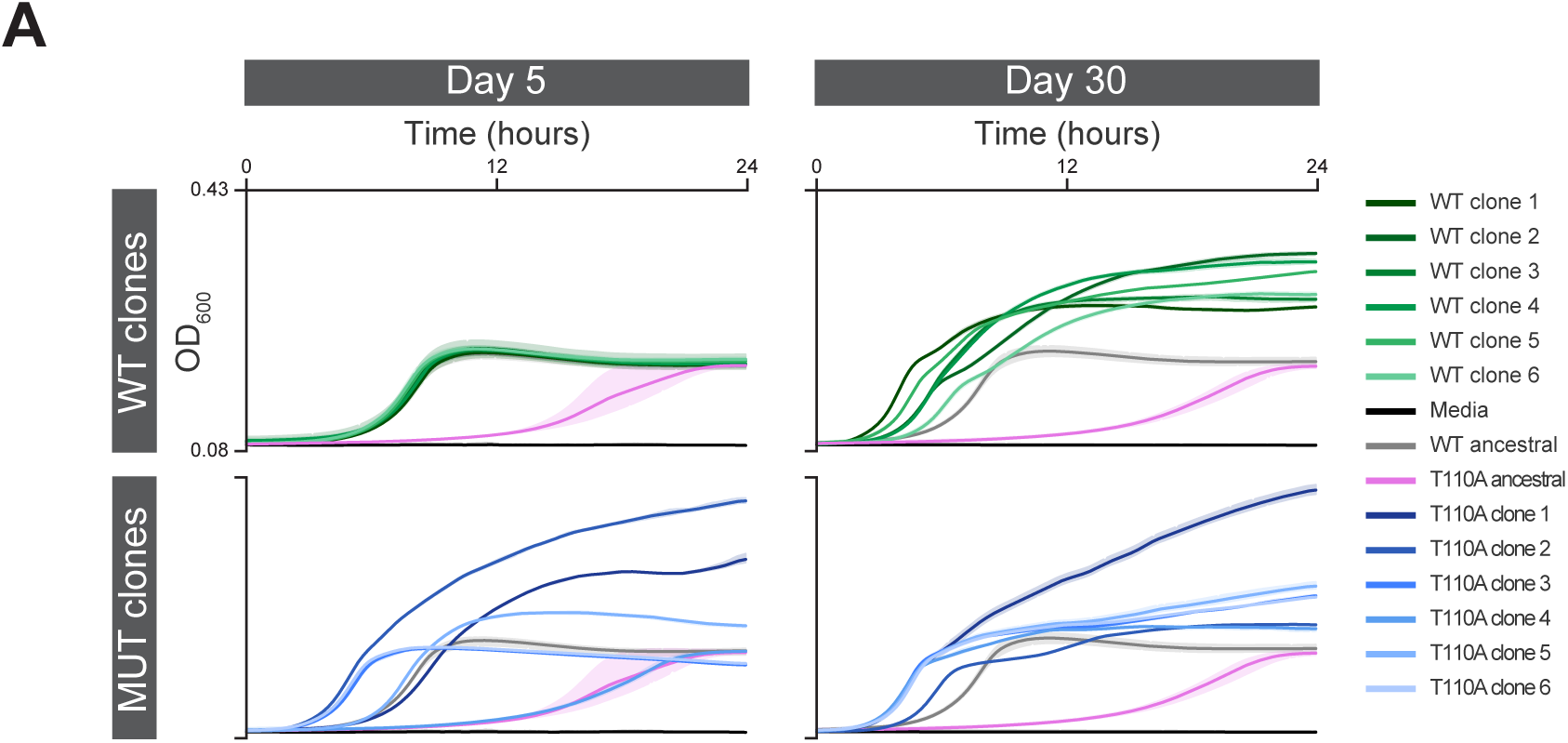
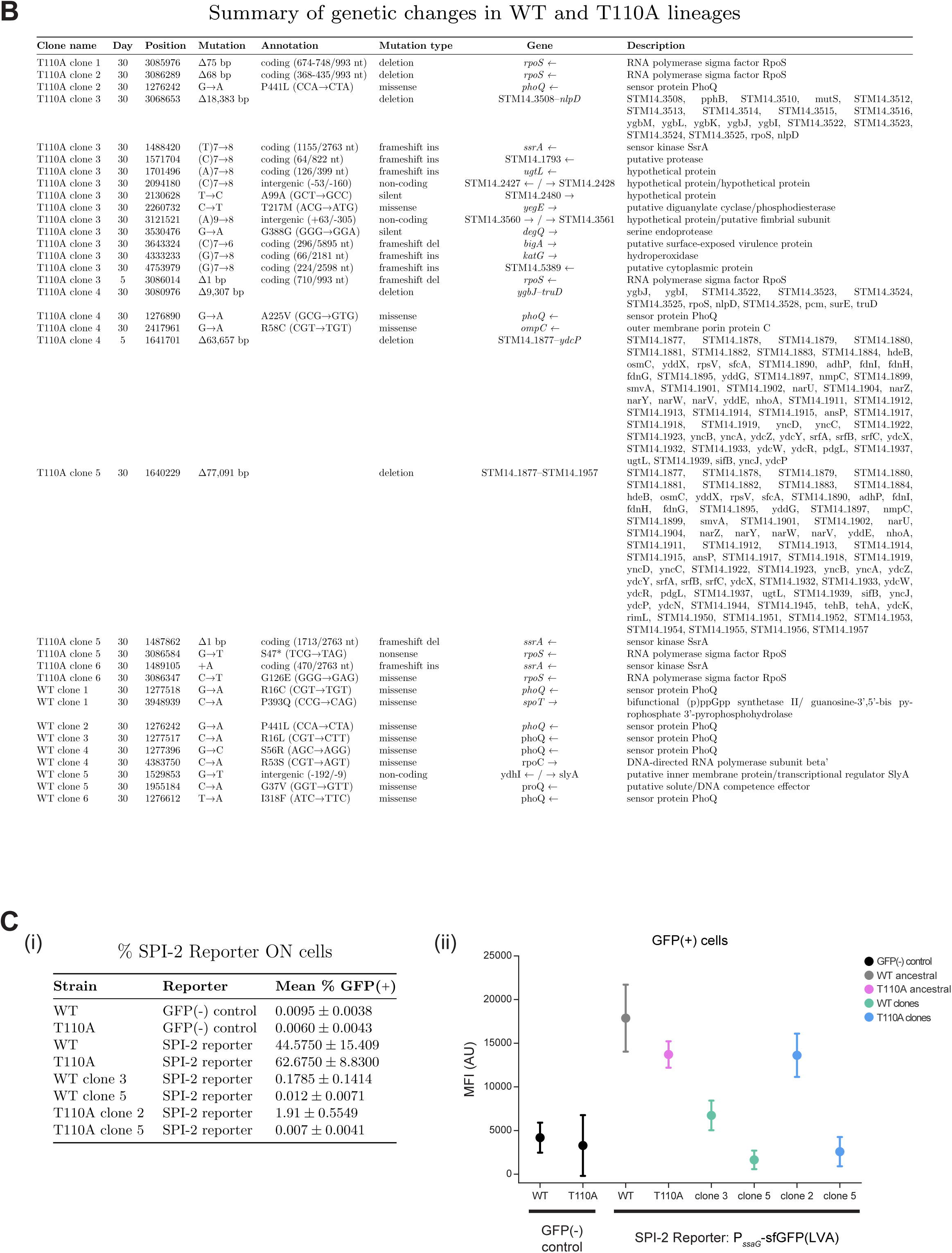
Growth, genotype, and SPI-2 reporter expression of evolved clones. (A) Biological replicate of growth curves for WT and T110A evolved clones at d5 and d30 (related to Figure 4A (ii)). (B) Summary of mutations in WT and T110A clones at d5 and d30. Three T110A clones (clones 1, 3, and 4) were sequenced at d5, while all six clones from each background were sequenced at d30. T110A clone 1 d5 had no mutations, but did have some new junction evidence as detailed in Table S1. (C) Loss of SPI-2 induction in evolved clones (related to Figure 4C). (i) Samples were analyzed by flow cytometry after 4 hours in MgM-MES and the percentage of GFP(+) cells is shown. Values represent mean ± standard deviation from four independent experiments. (ii) Mean fluorescence intensity (MFI) of GFP(+) cells. Only T110A clone 2 GFP(+) cells have GFP intensity comparable to the original WT and T110A strains. Other evolved clones show intensities similar to the GFP(−) control, indicating loss of SPI-2 induction. Data shown is mean ± standard deviation of 3 independent experiments. n=50,000 cells per sample for each replicate.

**Table S1.** Summary of whole genome sequencing results corresponding to data in Figures 1 and 4.

**Table S2.** Summary of data used for figures and statistical analyses.

**Table S3.** Reagents.

| REAGENT or RESOURCE | SOURCE | Catalog # |
| --- | --- | --- |
| **Salmonella Strains** |  |  |
| *Salmonella enterica* serovar Typhimurium 14028 WT | Michael McClelland Lab |  |
| *Salmonella enterica* serovar Typhimurium 14028 *hns* A328G (T110A) | This Study |  |
| **Salmonella Cell Culture** |  |  |
| LB broth, Miller | Fisher Scientific | BP1426 |
| LB broth, Lennox | Fisher Scientific | BP1427 |
| BactoAgar | Fisher Scientific | DF0140-01-0 |
| BD DIFCO™ Nutrient Agar | Fisher Scientific | DF0001-17-0 |
| BD Difco™ Dehydrated Culture Media: M9 Minimal Salts, 5x | Fisher Scientific | DF0485-17 |
| Calcium chloride dihydrate, for molecular biology, ≥99.0% | Sigma-Aldrich | C3306 |
| Magnesium sulfate heptahydrate, BioReagent, for molecular biology, suitable for plant cell culture, ≥99.0% | Sigma-Aldrich | M2773 |
| D-(+)-Glucose, powder, BioReagent, suitable for cell culture, suitable for insect cell culture, suitable for plant cell culture, ≥99.5% | Sigma-Aldrich | G7021 |
| Casamino Acids | Fisher Scientific | BP1424 |
| Potassium chloride, for molecular biology, ≥99.0% | Sigma-Aldrich | P9541 |
| Ammonium sulfate, for molecular biology, ≥99.0% | Sigma-Aldrich | A4418 |
| Potassium sulfate, BioXtra, ≥99.0% | Sigma-Aldrich | P9458 |
| Potassium phosphate monobasic, for molecular biology, ≥98.0% | Sigma-Aldrich | P9791 |
| Mes hydrate, BioPerformance Certified, suitable for cell culture, ≥99.5% | Sigma-Aldrich | M2933 |
| Glycerol | Fisher Scientific | BP229-1 |
| Magnesium chloride hexahydrate, BioReagent, suitable for cell culture, suitable for insect cell culture | Sigma-Aldrich | M2393 |
| **Cloning** |  |  |
| PrimeSTAR Max 2X Premix DNA Polymerase | Takara Bio | R045B |
| EcoFlex MoClo Kit | Addgene | Kit # 1000000080 |
| **Mammalian Cell Culture** |  |  |
| HeLa | ATCC | CCL-2 |
| Lenti-X 293T | Takara | 632180 |
| DMEM | Fisher Scientific | MT15017CV |
| Fetal Bovine Serum | Omega Scientific | FB-02 |
| Penicillin/Streptomycin | Fisher Scientific | 15-140-122 |
| GlutaMAX | Fisher Scientific | 35-050-061 |
| psPAX2 | Addgene | Plasmid #12260 |
| pMD2.g | Addgene | Plasmid #12259 |
| Hygromycin | Invivogen | ant-hg-1 |
| **Plate Reader** |  |  |
| Glass-bottom 96-well black plate | Tecan | 30122306 |
| AeraSeal film | Sigma-Aldrich | A9224 |
| **Whole Genome Sequencing** |  |  |
| Wizard(R) Genomic DNA Purification Kit | Promega | A1120 |
| **Protein Expression and Purification** |  |  |
| BL21(DE3) Competent Cells | Thermo Fisher Scientific | EC0114 |
| IPTG (Gold Biotechnology Inc) | Fisher Scientific | 50-153-2804 |
| Sodium chloride | Sigma-Aldrich | S5886-1KG |
| DL-Dithiothreitol, for molecular biology, ≥98% (HPLC), ≥99% (titration) | Sigma-Aldrich | D9779-5G |
| Phenylmethylsulfonyl Fluoride | Fisher Scientific | 52-332-5GM |
| cOmplete™, Mini, EDTA-free Protease Inhibitor Cocktail | Sigma-Aldrich | 11836170001 |
| Polyethyleneimine, approx. M.N. 60,000, 50 wt.% aq. solution, branched | Fisher Scientific | AC178571000 |
| 1-(3-Aminopropyl)imidazole 97.0+% | Fisher Scientific | A1185100G |
| 2-Mercaptoethanol, for molecular biology, for electrophoresis, suitable for cell culture, BioReagent, 99% (GC/titration) | Sigma-Aldrich | M3148-100ML |
| Invitrogen Ni-NTA Agarose | Fisher Scientific | R90101 |
| Sodium phosphate dibasic | Sigma-Aldrich | S3264-500G |
| Corning™ Cell Culture Phosphate Buffered Saline (1X) | Fisher Scientific | MT21040CV |
| PreScission Protease | Brickner Lab |  |
| Amicon Ultra Centrifugal Filter, 3 kDa MWCO | Sigma-Aldrich | UFC500396 |
| Glycine | Sigma-Aldrich | G8898-500G |
| Trizma® base | Sigma-Aldrich | 93362-250G |
| Sodium dodecyl sulfate | Sigma-Aldrich | L3771-500G |
| 2x Laemmli Sample Buffer | Bio-Rad | 1610737 |
| 4–15% Mini-PROTEAN® TGX™ Precast Protein Gels, 15-well, 15 µl | Bio-Rad | 4561086 |
| Precision Plus Protein™ Kaleidoscope™ Prestained Protein Standards | Bio-Rad | 1610375 |
| Bio-Safe Coomassie Stain | Bio-Rad | 1610786 |
| **EMSAs** |  |  |
| DNA Clean & Concentrator-5 | Zymo Research | D4004 |
| Bromophenol Blue | Fisher Scientific | AAA1846909 |
| Electrophoretic Mobility-Shift Assay (EMSA) Kit, with SYBR™ Green & SYPRO™ Ruby EMSA stains | Fisher Scientific | E33075 |
| 5% Mini-PROTEAN® TBE Gel, 15 well, 15 µl | Bio-Rad | 4565016 |
| Thermo Scientific™ LightShift™ Poly (dI-dC) | Fisher Scientific | PI20148E |
| **Flow Cytometry** |  |  |
| Electron Microscopy Sciences 16% Paraformaldehyde Aqueous Solution, EM Grade | Fisher Scientific | 50-980-487 |
| **Microscopy** |  |  |
| Fluorobrite DMEM | Fisher Scientific | A1896701 |
| Fibronectin | Sigma-Aldrich | F0895 |
| Cellvis 96 Well glass bottom plate with high performance #1.5 cover glass | Fisher Scientific | NC0536760 |
| Gibco™ Gentamicin (10 mg/mL) | Fisher Scientific | 15-710-064 |
| **Software and Algorithms** |  |  |
| Python3 | https://www.python.org/ |  |
| Seaborn | https://seaborn.pydata.org |  |
| Matplotlib | https://matplotlib.org/ |  |
| Cellpose3 | https://github.com/MouseLand/cellpose |  |
| CellTK | https://github.com/braysia/CellTK |  |
| Napari | https://napari.org/stable/ |  |
| nsbatwm | https://github.com/haesleinhuepf/napari-segment-blobs-and-things-with-membranes |  |
| clEsperanto | https://github.com/clEsperanto |  |
| Breseq | https://github.com/barricklab/breseq/ |  |
| FlowJo | v10.8.2 |  |
| Omniplate | https://git.ecdf.ed.ac.uk/pswain/omniplate |  |
| ChimeraX | https://www.rbvi.ucsf.edu/chimerax/ |  |
| Fiji | https://imagej.net/software/fiji/ |  |

**Table S4.** Plasmids and Bacterial Strains.

| **Plasmids** |  |  |  |
| --- | --- | --- | --- |
| **Plasmid Contents** | **Description** | **Antibiotic Resistance** | **Source** |
| pTU2-a-PssaG-RiboJ-Bba_B0064-sfGFP-LVA-L3S2P21-SJM901-PET_RBS-mRuby2-Bba_B0012-KanR-pSC101 | P<sub>ssaG</sub>-sfGFP-LVA (SPI-2 Reporter) | Kanamycin | Spratt *et al.* (2025) |
| pTU2-a-PinvF-RiboJ-Bba_B0064-sfGFP-L3S2P21-SJM901-PET_RBS-mRuby2-Bba_B0012-KanR-pSC101 | P<sub>invF</sub>-sfGFP (SPI-1 Reporter) | Kanamycin | This study |
| pTU2-a-PROMOTERLESS-sfGFP-L3S2P21-SJM901-PET_RBS-mRuby2-Bba_B0012-KanR-pSC101 | Promoterless Control Plasmid | Kanamycin | Spratt *et al.* (2025) |
| pTU1-B-SJM901-PET_RBS-mRuby2-Bba_B0012-AmpR-pSC101 | P<sub>SJM901</sub>-mRuby2 (Constitutive mRuby2) | Carbenicillin | Spratt *et al.* (2025) |
| pET-28a-H-NS-WT-PPX-6xHis | H-NS WT expression plasmid | Kanamycin | This study |
| pET-28a-H-NS-T110A-PPX-6xHis | H-NS T110A expression | Kanamycin | This study |

| **Bacterial Strains** |  |  |
| --- | --- | --- |
| **Strain contents** | **Description** | **Source** |
| STm 14028 WT P<sub>ssaG</sub>-sfGFP-LVA | WT with SPI-2 reporter plasmid | Spratt *et al.* (2025) |
| STm 14028 *hns* a328g P<sub>ssaG</sub>-sfGFP-LVA | T110A with SPI-2 reporter plasmid | This study |
| STm 14028 WT P<sub>invF</sub>-sfGFP | WT with SPI-1 reporter plasmid | This study |
| STm 14028 *hns* a328g P<sub>invF</sub>-sfGFP | T110A with SPI-1 reporter plasmid | This study |
| STm 14028 WT Promoterless Control Plasmid | WT with Promoterless control plasmid | Spratt *et al.* (2025) |
| STm 14028 *hns* a328g Promoterless Control Plasmid | T110A with Promoterless control plasmid | This study |
| STm 14028 WT P<sub>SJM901</sub>-mRuby2 | WT with Constitutive mRuby2 plasmid | Spratt *et al.* (2025) |
| STm 14028 *hns* a328g P<sub>SJM901</sub>-mRuby2 | T110A with Constitutive mRuby2 | This study |
| STm 14028 WT evolved clone 1 | WT evolved clone 1 Day 30 | This study |
| STm 14028 WT evolved clone 2 | WT evolved clone 2 Day 30 | This study |
| STm 14028 WT evolved clone 3 | WT evolved clone 3 Day 30 | This study |
| STm 14028 WT evolved clone 4 | WT evolved clone 4 Day 30 | This study |
| STm 14028 WT evolved clone 5 | WT evolved clone 5 Day 30 | This study |
| STm 14028 WT evolved clone 6 | WT evolved clone 6 Day 30 | This study |
| STm 14028 *hns* a328g evolved clone 1 | T110A evolved clone 1 Day 30 | This study |
| STm 14028 *hns* a328g evolved clone 2 | T110A evolved clone 2 Day 30 | This study |
| STm 14028 *hns* a328g evolved clone 3 | T110A evolved clone 3 Day 30 | This study |
| STm 14028 *hns* a328g evolved clone 4 | T110A evolved clone 4 Day 30 | This study |
| STm 14028 *hns* a328g evolved clone 5 | T110A evolved clone 5 Day 30 | This study |
| STm 14028 *hns* a328g evolved clone 6 | T110A evolved clone 6 Day 30 | This study |
| STm 14028 WT evolved clone 3 P<sub>ssaG</sub>-sfGFP-LVA | WT evolved clone 3 Day 30 with SPI-2 reporter plasmid | This study |
| STm 14028 WT evolved clone 5 P<sub>ssaG</sub>-sfGFP-LVA | WT evolved clone 5 Day 30 with SPI-2 reporter plasmid | This study |
| STm 14028 *hns* a328g evolved clone 2 P<sub>ssaG</sub>-sfGFP-LVA | T110A evolved clone 2 Day 30 with SPI-2 reporter plasmid | This study |
| STm 14028 *hns* a328g evolved clone 5 P<sub>ssaG</sub>-sfGFP-LVA | T110A evolved clone 5 Day 30 with SPI-2 reporter plasmid | This study |

